# A phage satellite tunes inducing phage gene expression using a domesticated endonuclease to balance inhibition and virion hijacking

**DOI:** 10.1101/2020.11.28.402263

**Authors:** Zoe Netter, Caroline M. Boyd, Tania V. Silvas, Kimberley D. Seed

## Abstract

Bacteria persist under constant threat of predation by bacterial viruses (phages). Bacteria-phage conflicts result in evolutionary arms races often driven by mobile genetic elements (MGEs). One such MGE, a phage satellite in *Vibrio cholerae* called PLE, provides specific and robust defense against a pervasive lytic phage, ICP1. The interplay between PLE and ICP1 has revealed strategies for molecular parasitism allowing PLE to hijack ICP1 processes in order to mobilize. Here, we describe the mechanism of PLE-mediated transcriptional manipulation of ICP1 structural gene transcription. PLE encodes a novel DNA binding protein, CapR, that represses ICP1’s capsid morphogenesis operon. Although CapR is sufficient for the degree of capsid repression achieved by PLE, its activity does not hinder the ICP1 lifecycle. We explore the consequences of repression of this operon, demonstrating that more stringent repression achieved through CRISPRi restricts both ICP1 and PLE. We also discover that PLE transduces in modified ICP1-like particles. Examination of CapR homologs led to the identification of a suite of ICP1-encoded homing endonucleases, providing a putative origin for the satellite-encoded repressor. This work unveils a facet of the delicate balance of satellite-mediated inhibition aimed at blocking phage production while successfully mobilizing in a phage-derived particle.

## Introduction

Mobile genetic elements (MGEs), gene cassettes capable of self-mobilization within and among genomes, play a powerful role in modulating interactions between bacteria and bacterial viruses (bacteriophages or phages). MGEs sit at the interface of bacteria-phage interactions by facilitating both the spread and evolution of anti-phage defense genes. Phage defense strategies, such as CRISPR-Cas systems (1), restriction modification systems (2), and other anti-phage systems are frequently encoded on MGEs (termed “defense islands”) (3) and are thus able to mobilize and spread. These systems and the genes they encode can protect a bacterial host from infecting phages, or conversely, allow infecting phages to overcome bacterial host defenses (4,5). In addition to facilitating mobilization, MGEs are a major driver of the evolutionary adaptation of defense genes and systems. Excision from or integration into a genome can cause regulatory alterations or mutations in existing genes, resulting in adaptive changes to existing defense genes or expression patterns (6–8). Genes delivered by MGEs can also be co-opted by the host for defense or unrelated core host functions via domestication, where mutations alter function in favor of the host (4,9,10).

The MGEs involved in transmitting defense systems display diverse self-proliferation mechanisms. Some are simply a single gene, such as homing endonuclease genes (HEGs). HEGs encode site-specific nucleases that cut DNA to stimulate host-mediated recombination-based repair, favoring insertion of the HEG sequence into the target site (11). Other MGEs like plasmids and transposons require a combination of genes and noncoding DNA features (e.g. origins of replication and inverted repeat sequences) for replication and transfer (12). The most complex MGEs are phages themselves, which encode structural components to package their genomes and arrays of host-takeover and counter-defense genes. An intriguing and complex class of sub-viral MGEs are phage satellites, which lie dormant in a host genome and rely on helper phages for activation and packaging components in order to proliferate. Lytic infection by a helper phage stimulates satellite excision, genome replication and gene expression which redirects phage packaging to favor satellite genome transmission via transduction in phage-derived particles (13,14).

*Vibrio cholerae*, the bacterial causative agent of the disease cholera, can encode remarkable phage satellites called PLEs (phage-inducible chromosomal island-like elements) (15). PLEs display canonical excision, replication, and self-transmission activity in a *V. cholerae* host infected with an environmentally dominant lytic vibriophage, ICP1 (15–18). While many mechanisms of PLE activity remain enigmatic, PLEs effectively protect *V. cholerae* populations from lytic ICP1 predation by a suite of strategies including inhibiting ICP1 genome replication (17) and causing accelerated lysis of the infected *V. cholerae* host population, which interrupts ICP1 progeny phage construction (15,18).

Mechanistic studies of PLE activity have revealed many functional similarities to the well-characterized phage satellites (the *<u>S</u>taphylococcus <u>a</u>ureus* <u>p</u>athogenicity islands, or SaPIs), as well as distinct layers of complexity that highlight PLE’s specific co-evolution with and dependence on ICP1 (Supplementary Figure S1) (16–18). Understanding PLE-mediated processes can help define mechanistic paradigms of the poorly-understood family of phage satellites and perhaps uncover novel innovations adapted specifically by PLE. There is much to be discovered regarding the later stages of PLE activity, including packaging and transduction. SaPIs, along with other viral satellites, are known to exploit their helper virus’s structural components to construct their own transducing particles (13). SaPIs have smaller genomes than their inducing phage and package their DNA into phage-derived particles with reduced capsid diameter. Because the larger phage genome cannot be completely packaged into a SaPI-sized capsid with a smaller diameter, this mechanism effectively reduces viable phage output from an infection with an active SaPI. Thus, in SaPIs, this redirection of capsid size serves as a direct phage interference mechanism (19). While PLEs have been shown to transduce with the same receptor requirement as ICP1, they transduce at a low frequency compared to SaPIs (15), and the morphology and constitution of PLE transducing particles is currently unknown.

Despite their disparate origins, hosts, and sequence content, PLEs and certain SaPIs both repress their helper phage’s structural gene transcription. Some SaPIs encode *ptiA*, a robust inhibitor of helper phage late gene transcription that interacts with the phage’s late transcriptional activator LtrC (20). Recent global transcriptomic analysis of ICP1 infection of *V. cholerae* in the presence and absence of PLEs revealed hints of a similar but more targeted PLE-mediated impact on ICP1 transcription (21). We observed that a single operon in the ICP1 genome is transcriptionally downregulated in a PLE(+) host. The three-fold downregulated operon (*gp126-gp122*, from here forward referred to as the capsid operon) encodes ICP1’s capsid morphogenesis proteins including the major capsid subunit (Gp122), putative capsid decoration protein (Gp123), putative capsid maturation protease (Gp125) and other proteins with unknown function (Figure 1A) (21). This subtle and specific impact on ICP1 transcription was surprising given that PLE activity during ICP1 infection restricts ICP1 DNA replication and completely eliminates production of phage progeny (15,17). Capsid operon repression is a conserved feature of PLEs: five PLEs with unique allelic variations identified in *V. cholerae* isolates all reduce transcript and protein produced from the capsid operon (21). This work will refer to the modern dominant PLE1 as “PLE” from here forward unless otherwise specified.

**Figure 1.**
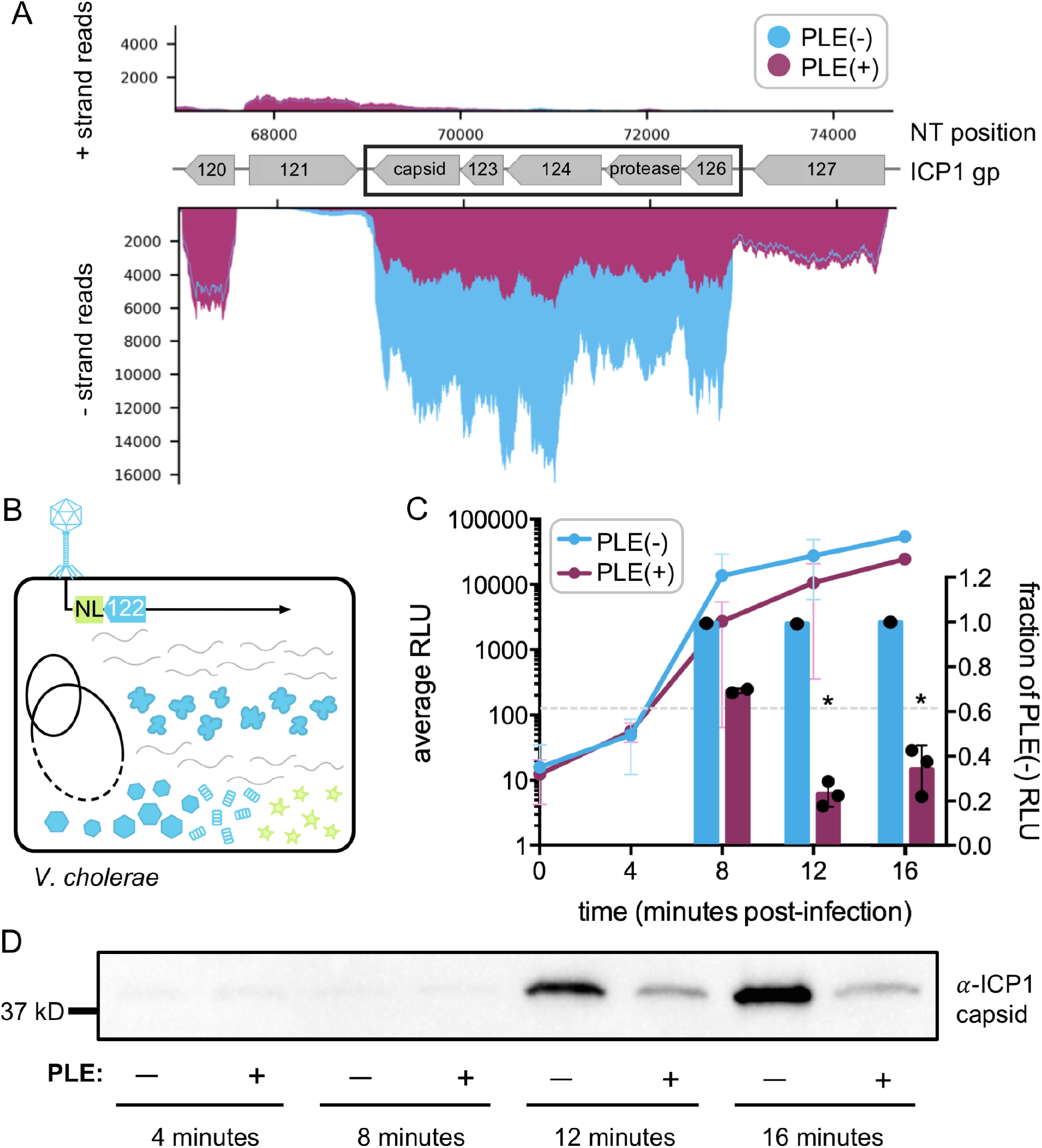
PLE specifically represses expression of ICPl’s capsid operon. **(A)** ICPl RNA­ sequencing reads (Ref. 21) comparing read coverage across the differentially expressed ICPl capsid operon (black rectangle) 16 minutes post-infection in a PLE(−) and a PLE(+) *V. cholerae* host. Top x-axis indicates ICPl genome nucleotide (NT) position and grey gene arrows beneath indicate the predicted open reading frame number/predicted function. **(B)** Schematic of **ICPl** capsid operon reporter phage used throughout this study. NanoLuc **(NL)** luminescent reporter (yellow star) is expressed under the native operon promoter immediately downstream of the last gene in the operon *(gpl 22*, the capsid monomer). (C) Left - line graph plots raw luminescent output at the timepoints indicated post-infection with the ICPl capsid operon reporter phage in PLE(−) and PLE(+) *V. cholerae* host strains. Right - bar graphs express the raw luminescence as a fraction of the luminescent output of a PLE(−) infection. Grey dashed line indicates RLU limit of detection. *p < 0.0005 Student t-test by column pair (n=3). (D) Representative Western blot against ICPl capsid (direct peptide antibody) over ICPl infection time-course in PLE(−) and PLE(+) *V. cholerae* hosts. Total protein from cell lysate at each timepoint was calculated and normalized to standardize total protein input into each lane.

An expectation for more robust global transcriptional downregulation of ICP1 by PLE was also founded in the observation that immediately prior to lysis during a PLE(+) ICP1 infection, PLE genome copy outnumbers ICP1 genome copy by eight-fold late in infection, suggesting that ICP1 DNA available as a template for late gene transcription is significantly reduced by PLE DNA replication (17). The precise impact on a single operon in late ICP1 transcription is not obviously congruent with PLE’s global impact on ICP1 DNA replication, so we first sought to develop new tools to probe precise expression levels at specific ICP1 loci. We utilize these tools to uncover the PLE-encoded mechanism responsible for transcriptional repression of the ICP1 capsid operon: CapR, a DNA-binding protein that we hypothesize is derived from a phage homing endonuclease. This instance of HEG co-option for a highly conserved satellite function suggests that such genetic exchange can provide important substrates for phage-satellite co-evolution.

## Materials & Methods

### Biological Resources

All strains used in this study are listed in Supplementary Table S2. *V. cholerae* mutant strains were constructed via natural transformation of linear PCR products synthesized by SOE (splicing by overlap extension) PCR (22). Primer sequences available upon request. ICP1 reporter phage and CRISPRi strains were constructed via CRISPR-based editing system as previously described (23) by introducing an editing template containing the mutation or nanoluciferase open reading frame in the appropriate genetic context (for nanoluc reporters: immediately downstream of ICP1 *gp68* or *gp122*).

### Bacterial & phage growth conditions

*V. cholerae* strains were maintained in 20% glycerol stocks stored at −80°C and propagated at 37°C by streaking onto LB agar plates and inoculation into Miller LB liquid media (Fisher Scientific) with aeration. Antibiotics were added to media at the following concentrations where appropriate: kanamycin (75μg/mL), spectinomycin (100μg/mL), and ampicillin (50μg/mL). For cultures with inducible *capR* expression or chromosomal empty vector, 1mM isopropyl β-D-1-thiogalactopyranoside (IPTG) and 1.5mM theophylline were added to induce expression of the construct upon back-dilution prior to infection. For CRISPRi induction, 1mM isopropyl β-D-1-thiogalactopyranoside (IPTG) was added upon back-dilution to OD_600_=0.05 prior to infection. ICP1 phage strains were propagated on *V. cholerae* hosts by soft agar overlay method (0.5% agar in LB, 37°C for 6 hours). To prepare high-titer stocks, phages were harvested after overnight rocking at 4°C from an overlay of STE buffer (1M NaCl, 200mM Tris-HCl, 100mM EDTA) on confluent lysis plates. Phage lysate was collected, chloroform-treated, and cleared to remove bacterial debris before concentrating phage to generate high titer stocks by polyethylene glycol precipitation (24).

For phage infections, *V. cholerae* host strains were grown with appropriate antibiotics to OD_600_ > 1 in liquid culture at 37°C, then back-diluted to OD_600_ = 0.05 before growing up to OD_600_ = 0.3 to infect with phage at the appropriate multiplicity of infection (MOI).

### Nanoluc reporter phage assays

*V. cholerae* host strains were grown to OD_600_ = 0.3 from a culture grown to OD_600_ > 1 then back-diluted to OD_600_ = 0.05. These cultures were infected at an MOI = 0.1 with ICP1 reporter phage encoding the NanoLuc nanoluciferase gene (25) and returned to incubate at 37°C with aeration. At each time-point, a 100μl sample was taken from each infection and mixed with an equal volume of ice-cold methanol. Larger numbers of strains were screened with ICP1 reporter phage by infecting multiple 200μl aliquots of each *V. cholerae* strain in clear 96-well plates and harvesting from new wells for each timepoint with a multichannel pipette. Luminescence was measured from the samples with the Nano-Glo Luciferase Assay System (Promega) according to manufacturer protocol with a SpectraMax i3x microplate reader (Molecular Devices). Samples were aliquoted on ice in every other well of black half-area 96-well plates (Corning) to avoid signal bleed-through and measured every minute for 6 minutes-post substrate addition to capture an average signal. Experiments were conducted in biological triplicate.

### Western blots

*V. cholerae* host strains were grown to OD_600_ = 0.3 as described above. Cultures were infected with phage at an MOI = 1 and returned to incubate at 37°C with aeration. At each time-point, a 1mL sample was collected from the infection and mixed with an equal volume of ice-cold methanol. Samples were centrifuged at 5,000xg for 10 minutes at 4°C to pellet. Pellets were washed once with ice-cold PBS and resuspended in cold lysis buffer (50mM Tris, 150mM NaCl, 1mM EDTA, 0.5% Triton X-100, 1x Pierce™ Protease Inhibitor Mini Tablet (Thermo)). Protein concentration was quantified with Pierce BCA Protein Assay Kit (Thermo), and 30μg of total protein per sample was mixed with Laemmli buffer supplemented with 2-mercaptoethanol (10%) (Bio-rad) and boiled at 99°C for 10 minutes. Samples were run on Any-kD TGX-SDS-PAGE gels (Bio-rad) and transferred onto nitrocellulose membranes with Transblot Turbo Transfer system (Bio-rad). Custom primary peptide antibody generated in rabbits against ICP1 capsid (GenScript) was diluted 1:1500 and applied to the membrane for more than three hours. Band detection was conducted with a goat α-rabbit-HRP secondary antibody (Bio-rad) at 1:10,000 followed by development with Clarity Western ECL Substrate (Bio-rad) and imaging on a Chemidoc XRS Imaging System (Bio-rad). Band quantification from biological triplicate experiments was conducted using ImageJ [https://imagej.nih.gov/ij/].

### Transduction assays

*V. cholerae* strains with either a wild-type or Δ*capR* PLE marked with a kanamycin resistance cassette downstream of *orf23* (as in (15)) were grown to OD_600_ = 0.3 and infected with ICP1 at an MOI = 2.5. Following 5 minutes at 37°C for attachment, the infected cultures were then washed twice in 1 volume warm LB to remove excess phage, then resuspended in LB (or LB supplemented with 10mM MgCl_2_ for direct comparison). Cultures were incubated for 30 minutes at 37°C with aeration then treated with chloroform and centrifuged at 4,000xg for 10 minutes to remove debris. Cleared lysate was added at a 1:10 ratio to an overnight culture of a PLE(−) spectinomycin-resistant recipient strain (Δ*lacZ:*spec) supplemented with 10mM MgCl_2_ after overnight growth. Recipients were mixed with transducing lysate for 20 minutes at 37°C with aeration before plating on LB plates supplemented with kanamycin (75μg/mL) and spectinomycin (100ug/mL) to select for transductants in technical duplicate. Assay was performed in biological triplicate.

### ICP1 one-step burst assays

*V. cholerae* strains (see Supplementary Table S2 for relevant genotypes) were grown to OD_600_ = 0.3 in LB supplemented with inducer (see Bacterial growth conditions) and infected with ICP1 at an MOI = 0.1. After 5 minutes of attachment at 37°C, infected cultures were diluted 1:2500 then serially 1:10 and returned to incubate at 37°C. At each listed time-point, a sample was harvested and lysed with chloroform. Phage titer at each time-point was determined in technical triplicate by plaque assay. Assay was performed in biological triplicate.

### Protein purification

*E. coli* BL21 cells were transformed with a pE-6xHis-SUMO-CapR fusion construct and grown in Terrific Broth supplemented with kanamycin (75μg/mL) at 37°C to an OD_600_=0.6. Cultures were then induced with addition of 0.5mM isopropyl β-D-1-thiogalactopyranoside (IPTG) and incubated overnight at 16°C with aeration. Cells were collected by centrifugation and resuspended in lysis buffer (50mM NaH_2_PO_4_, 150mM NaCl, 20mM imidazole, 1% glycerol, 2mM 2-mercaptoethanol, 0.5% Triton X-100, 1x Pierce™ Protease Inhibitor Mini Tablet (Thermo), pH 7.5) prior to sonication at 4°C. Lysate was cleared by centrifugation (20,000xg, 30 minutes 4°C), treated with DNase and RNase (100μl each) on ice for 20 minutes, and filtered through 0.2μm cellulose filters (GE Life Sciences). Cleared lysate was applied to a pre-equilibrated HisTrap (Sigma) Ni-sepharose column and washed with 10 column volumes wash buffer I (50mM NaH_2_PO_4_, 150mM NaCl, 70mM imidazole, 1% glycerol, 2mM 2-mercaptoethanol, pH 7.5) followed by 5 column volumes high salt wash buffer II (50mM NaH_2_PO_4_, 2.5M NaCl, 20mM imidazole, 1% glycerol, 2mM 2-mercaptoethanol, pH 7.5), followed by 7 column volumes base buffer (50mM NaH_2_PO_4_, 150mM NaCl, 20mM imidazole, 1% glycerol, 2mM 2-mercaptoethanol, pH 7.5). Protein was eluted in several small aliquots with imidazole elution buffer (50mM NaH_2_PO_4_, 150mM NaCl, 250mM imidazole, 1% glycerol, 2mM 2-mercaptoethanol, pH 7.5). Glycerol was added to 20% final concentration for storage at −80°C. *E. coli* protein contaminants in the purified preparation used for *in vitro* assays were identified by 1D mass spectrometry performed from a gel slice to ensure no known DNA binding proteins co-purified with CapR at significant concentration (Supplementary Figure S9B).

### DNA binding and electrophoretic mobility shift assays (EMSAs)

DNA probes were synthesized via PCR with Q5 high-fidelity polymerase (Thermo) and purified with Monarch PCR Cleanup Kit (NEB). 75ng probe A/probe B and 150ng probe C/probe D were added to increasing concentrations of protein (4%, 25%, and 50% of reaction mix with remaining volume matched with addition of imidazole elution buffer (50mM NaH_2_PO_4_, 150mM NaCl, 250mM imidazole, 1% glycerol, 2mM 2-mercaptoethanol, pH 7.5)) in EMSA reaction buffer (50mM NaH_2_PO_4_, 150mM NaCl, 5% glycerol, 1mM dithiothreitol (DTT), 5mM MgCl_2_, pH 7.5). Reactions incubated for 25 minutes at 30°C before mixing gently with DNA loading dye before loading the entire reaction into a 5% polyacrylamide-Tris-Borate gel and running in 0.5X TB Buffer (45mM Tris, 45mM Boric Acid) for 45 minutes at 4°C. Gels were post-stained for 10 minutes in 0.5X TB Buffer with GelGreen (Biotium) and rinsed with sterile water before imaging with UV on an EZ Dock Imager (Bio-Rad). EMSAs were performed in biological triplicate (Supplementary Figure S9A contains additional replicates). Probe nucleotide positions in ICP1_2006E_ΔCRISPR/Cas genome: Probe A: 72881-72984, Probe B: 101037-101133, Probe C: 72881-72946, Probe D: 72922-72984

### Transmission electron microscopy (TEM)

Lysates were treated with chloroform and the cellular debris removed by centrifugation (5000xg at 4°C for 15 minutes). From the supernatant containing virions, a 5μL sample was applied to a copper mesh grid (Formvar/Carbon 300, Electron Microscopy Sciences) for 60 seconds, washed with sterile ddH_2_O for 15 seconds, and stained with 1% uranyl acetate for 30 seconds. Micrographs were collected with a FEI Tecnai-12 electron microscope operating at 120 kV. Multiple biological replicates were imaged for each ICP1 and PLE genotype. Measurements were taken from greater than or equal to eight particles prepared in parallel.

### Protein alignments

Pairwise protein alignments were conducted directly with MUSCLE (default parameters) [https://www.ebi.ac.uk/Tools/msa/muscle/] and visual estimations of pairwise alignments were constructed by hand. Multiple sequence alignments were conducted with MUSCLE (default parameters) and visualized with JalView [https://www.jalview.org/] or conducted and visualized with PRALINE (default parameters) [https://www.ibi.vu.nl/programs/pralinewww/].

### Statistical Analysis

All experiments were conducted in technical duplicate and biological triplicate (n=3) unless otherwise specified. p-values represent two-sided t-tests comparing two groups, replicates unpaired when comparing ratios and paired when comparing raw values. Error bars represent standard deviation.

## Results

### PLE represses ICP1 capsid protein expression

To validate and characterize the PLE-mediated ICP1 capsid operon repression detected by transcriptomics, we required a new reporter tool to assay transcriptional activity. This requirement was dictated by the unique challenges presented by the lifecycle of ICP1 within a *V. cholerae* host cell, which is rapid and highly destructive: infections produce ~90 viable phage progeny per infected cell in under 20 minutes (15). The bacterial host chromosome is rapidly degraded (26) and transcriptional profiling during infection did not reveal any ICP1, PLE, or *V. cholerae* genes with consistent transcriptional activity over the infection time-course (21). This limits feasibility of evaluating the targeted repression of the capsid operon by RT-qPCR, which would require a proper normalization control. Instead, we developed an *in vivo* reporter system to monitor temporal ICP1 gene expression during infection and enable subsequent genetic dissection of PLE-mediated targeted transcriptional manipulation. This reporter system harnesses the small, stable, and rapidly folding reporter protein NanoLuc nanoluciferase (NanoLuc) (25). Using CRISPR-Cas engineering (23) we constructed ICP1 reporter phages expressing NanoLuc downstream of genes of interest. We constructed an ICP1 reporter phage with a NanoLuc reporter integrated downstream of an ICP1 operon that is categorized by transcriptomics as expressed as a middle gene (*gp68*) as well as a phage strain with a reporter integrated downstream of the late-expressed *gp122* capsid gene, which is part of the differentially regulated ICP1 capsid operon (Figure 1B, Supplementary Figure S2A). Each of these phage strains were separately used to infect *V. cholerae* at a multiplicity of infection (MOI) of 0.1 and luminescence was monitored over the ICP1 infection cycle. The ICP1 middle gene (*gp68*) reporter phage unreliably produces luminescence above the limit of detection at 4 minutes post-infection, increases rapidly then plateaus between 12 and 16 minutes post-infection (Supplementary Figure S2B), while the late gene (*gp122*-*capsid*) reporter phage first produces detectable luminescent signal at 8 minutes post-infection and produces increasing luminescent signal over the duration of the infection (Figure 1C). The observed temporal reporter expression correlates with our transcriptomic observations, indicating that the *in vivo* reporter system adequately captures fine-scale temporal dynamics of ICP1 gene expression.

The capsid operon late-expressed reporter phage was used to infect both PLE(−) and PLE(+) *V. cholerae* hosts to assess if differential expression of this operon could be detected in the presence of PLE with the *in vivo* reporter system. In line with the three-fold decrease in capsid operon expression observed by transcriptomics (21), we observed that the capsid operon reporter phage in a PLE(+) infection produced approximately one-third of the reporter signal of a PLE(−) infection at 12 and 16 minutes post-infection (Figure 1C). To ensure the transcriptional decrease observed with the capsid operon reporter phage was specific to this late-expressed operon, we used the middle *gp68* reporter phage to infect PLE(−) and PLE(+) *V. cholerae* hosts and observed no difference in NanoLuc expression between hosts at any timepoint (Supplementary Figure S2B). This confirms that NanoLuc expression from the reporter phage can capture subtle differences in gene expression at specific loci.

To better define the impact of PLE-mediated ICP1 transcriptional repression at the capsid operon, we compared the amount of ICP1 capsid protein produced in PLE(+) and PLE(−) infections over time. Western blot analysis of total protein from PLE(−) and PLE(+) hosts over the course of infection demonstrated detectable capsid protein by 12 minutes post-infection and confirmed that PLE(+) infections produce approximately one-third as much ICP1 capsid protein as PLE(−) infections at all timepoints where capsid protein was detected (Figure 1D, Supplementary Figure S3A). These results together with the capsid operon reporter phage validate the observation of capsid operon transcriptional repression mediated by PLE (21) and confirm that this transcriptional repression results in a reduction in total ICP1 capsid monomer production during the late stages of a PLE(+) infection.

### A single PLE gene product is responsible for ICP1 capsid repression

We next sought to determine if a PLE-encoded gene product was necessary for mediating repression of ICP1’s capsid operon. Transcriptomics experiments revealed that all five previously described PLEs specifically targeted the ICP1 capsid operon resulting in similar degrees of transcriptional repression (21), indicating that one or more of the 11 core genes conserved in all five PLEs (15) was likely involved in this process. To test this, 11 PLE(+) strains each with a single core gene knocked out were infected with the ICP1 capsid operon reporter phage and luminescence was assessed as a fraction of the luminescence produced from a PLE(−) infection (Supplementary Figure S4). Deletion of *orf2* (from here forward referred to as *capR*, for capsid repressor) uniquely supported robust capsid operon expression during ICP1 infection on par with a PLE(−) infection (Figure 2A). *capR* was also necessary for PLE mediated repression of ICP1 capsid at the protein level as we observed a three to four-fold increase in ICP1 capsid by Western blot in PLE Δ*capR* compared to wild-type PLE (Figure 2B, Supplementary Figure S3B). The relief of capsid repression is not due to the necessity of *capR* for overall PLE function because previous analyses found no defect for a *capR* deletion in PLE excision (16), replication (17), PLE-mediated lysis, or PLE-mediated inhibition of ICP1 plaque formation (18).

**Figure 2.**
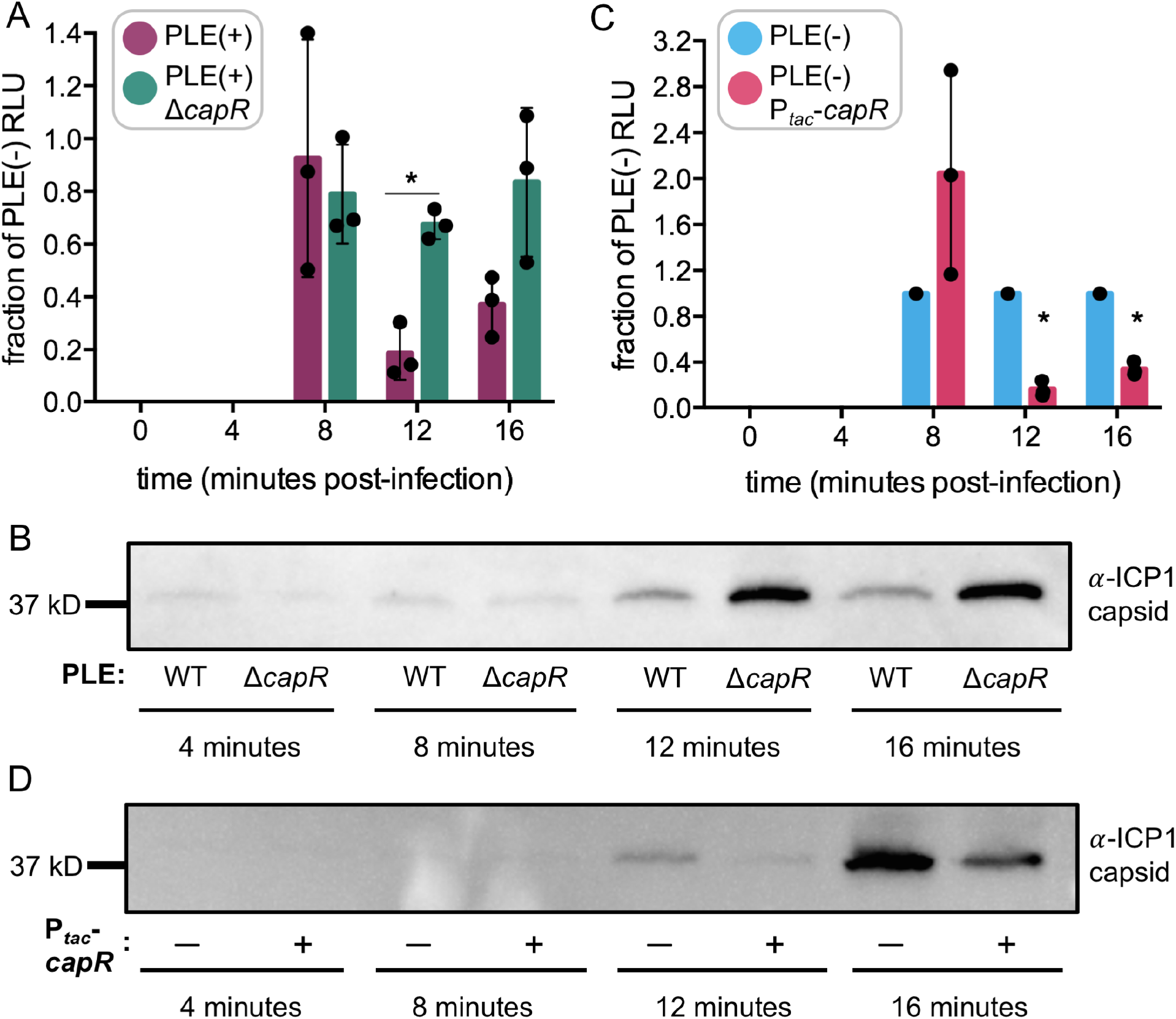
PLE CapR mediates ICPl capsid repression. **(A)** ICPl capsid operon reporter luminescence expressed as a fraction of luminescence from a PLE(−) reporter infection comparing wild-type PLE with a kanamycin resistance cassette integrated downstream of *orj23* (PLE(+)) and an isogenic strain with the *capR* open reading frame deleted *(!icapR).* *p < 0.002 by Student t-test per timepoint (n=3). (B) Western blot against ICPl capsid protein over ICPl infection time-course in *V. cholerae* host strains with wild-type PLE and PLE lacking *capR.* Total protein from cell lysate at each timepoint was calculated and normalized to standardize total protein input into each lane. (C) ICPl capsid operon reporter during infection in a host pre-induced to express *capR* from a cassette integrated in the *lacZ* locus of the *V. cholerae* chromosome *(P_iac_-capR)* (fraction of luminescence from a PLE(−) infection). *p<0.005 by Student t-test per timepoint (n=3) (D) Western blot against ICPl capsid over infection time-course with strains in (C). Total protein from cell lysate at each timepoint was calculated and normalized to standardize total protein input into each lane.

We next directly assessed if *capR* was sufficient to mediate repression of ICP1’s capsid operon outside the context of PLE. *capR* expressed from an inducible cassette integrated in the *lacZ* locus in PLE(−) *V. cholerae* recapitulated repression of ICP1’s capsid to approximately one-third of reporter expression in a PLE(−) empty vector control (Figure 2C) and by anti-ICP1 capsid Western blot of an infection time-course (Figure 2D, Supplementary Figure S3C). The capsid depletion observed by Western blot is not due to *capR*-mediated toxicity that could broadly impact phage infection dynamics, as the CapR expression strain grew normally with induction (Supplementary Figure S5A). These results demonstrate that *capR* is sufficient to mediate repression of ICP1’s capsid operon to the same degree as observed with total PLE activity.

The *capR* gene product has a subtle effect on ICP1 transcription in the context of PLE, so we were interested to see if induced expression of this gene would be sufficient to interfere with ICP1 progeny production. CapR reduces the amount of ICP1 capsid monomer (and other genes we expect to be essential phage structural components based on their inclusion in the operon), so we suspected that expression of *capR* might reduce the amount of ICP1 progeny from an infection by reducing the number of monomers available to form complete phage capsid heads, which typically require hundreds of monomers each to assemble. Somewhat surprisingly, expression of *capR* did not reduce ICP1 efficiency of plaquing (Figure 3A, Supplementary Figure S5B). This assay enumerates the number of hosts that were successfully infected and supported enough lytic phage infection to form a visible clearing (plaque), so its sensitivity to more subtle differences in a single round of infection is limited. In order to quantify the ICP1 progeny produced in a single round of infection with *capR* expression, we performed one-step burst assays with the inducible *capR* expression strain and saw no significant difference between the number of ICP1 progeny particles produced with or without CapR (Figure 3B). Because CapR activity reduces the expression of the ICP1 capsid operon but does not completely repress it, we take this data to suggest that the degree of capsid operon expression allowed under CapR-mediated repression is sufficient for ICP1 replication under the conditions tested. Since we did not find evidence that CapR activity is directly antagonistic to ICP1, we hypothesized that it may instead be playing an important role in a conserved PLE activity like transduction.

**Figure 3.**
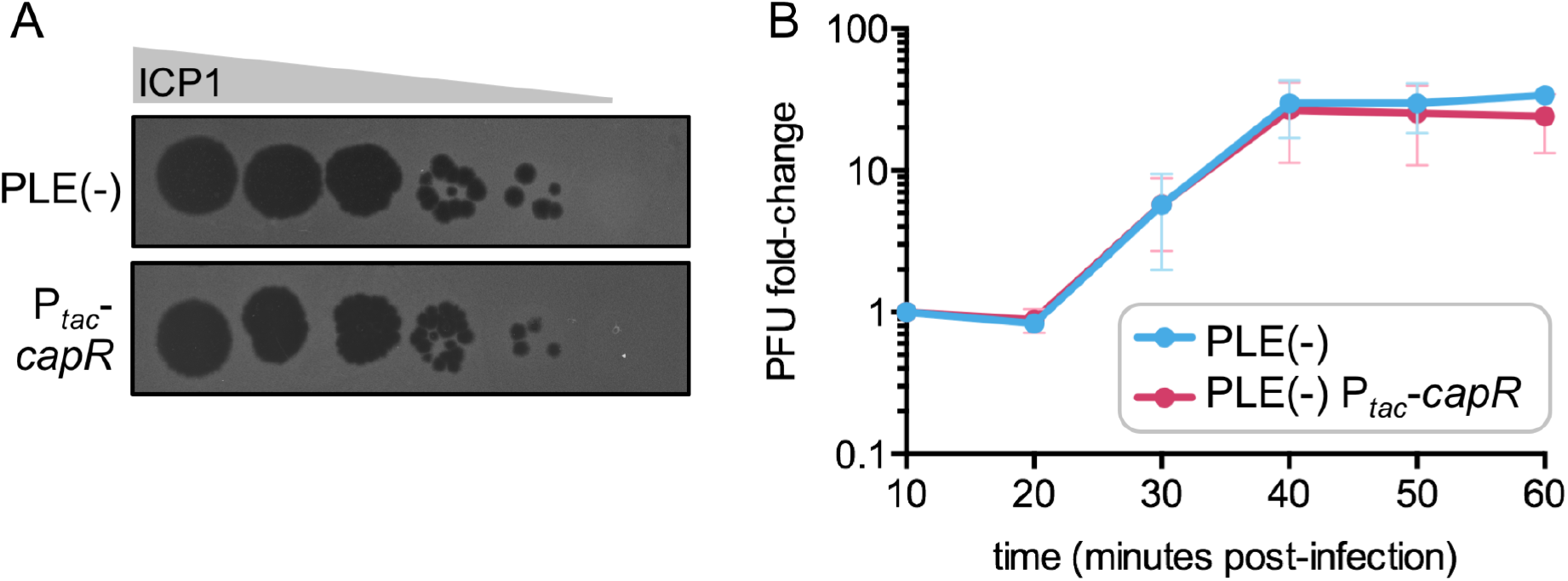
CapR expression alone does not interfere with ICPl progeny production. (A) ICPl phage spot assays. Lytic ICPl phage form plaques (clearings, black spots) on a V *cholerae* host lawn (grey) over multiple rounds of phage replication. Spots represent~ 10-fold serial dilutions oflCPl. PLE(−) host lawn (grey) expresses an empty vector (PLE(−)) or *capR* (P,ac*-capR)* from an inducible cassette integrated in the *lacZ* locus of the *V cholerae* chromosome. **(B)** ICPl one-step burst assay quantifying the number of ICPl plaque forming units (PFU) produced over a single round of infection in empty vector and capR-expressing hosts.

PLE replicates its genome ~1000-fold during ICP1 infection (15). PLE genome replication plays a role in ICP1 restriction (17) and also suggests that each PLE(+) cell could theoretically produce up to 1000 transducing particles. However, PLE transduction efficiency was previously reported to be very low (less than one functional transducing unit per thousand infected PLE1(+) donor cells) (15). PLE also does not encode any genes predicted to function in capsid morphogenesis, suggesting that like other phage satellites, PLE parasitizes ICP1-encoded virion proteins for transduction. We therefore hypothesized that PLE repurposes ICP1 structural components for its own transfer and that these restructured particles could be less stable than ICP1 virions or have stricter host-membrane integrity requirements than ICP1 for successful attachment. Enhanced phage particle stability can be achieved with addition of divalent magnesium cations (Mg^2+^) (27). Similarly, the Gram-negative bacterial outer membrane consists of a layer of lipopolysaccharide molecules, each containing negatively-charged phosphate groups in the core region that can be neutralized and bridged by cations including Mg^2+^ (28,29) Magnesium appears to play a role in both phage particle and host membrane stability, therefore we evaluated if supplemental magnesium cations could elevate levels of PLE transduction. Strikingly, we observed that with addition of 10mM MgCl_2_, PLE transduction efficiency was enhanced by two orders of magnitude (Figure 4A). Previous efforts to characterize PLE transducing particles were hindered by the apparently low level of transduction, but with increased transduction efficiency we were able to evaluate the morphology of PLE transducing particles. Electron microscopy performed on lysates produced from ICP1 infection of PLE(+) *V. cholerae* in the presence of magnesium revealed morphologically distinct PLE transducing particles with smaller heads measuring approximately 50nm in diameter, compared to 80nm diameter ICP1 capsid heads (Figure 4E). PLE and ICP1 heads have the same icosahedral shape, and PLE transducing particle tails appear to have the same morphology as ICP1 tails (Figure 4B). This morphological evidence further supports PLE’s piracy of ICP1 structural components for its own transducing particle assembly.

**Figure 4.**
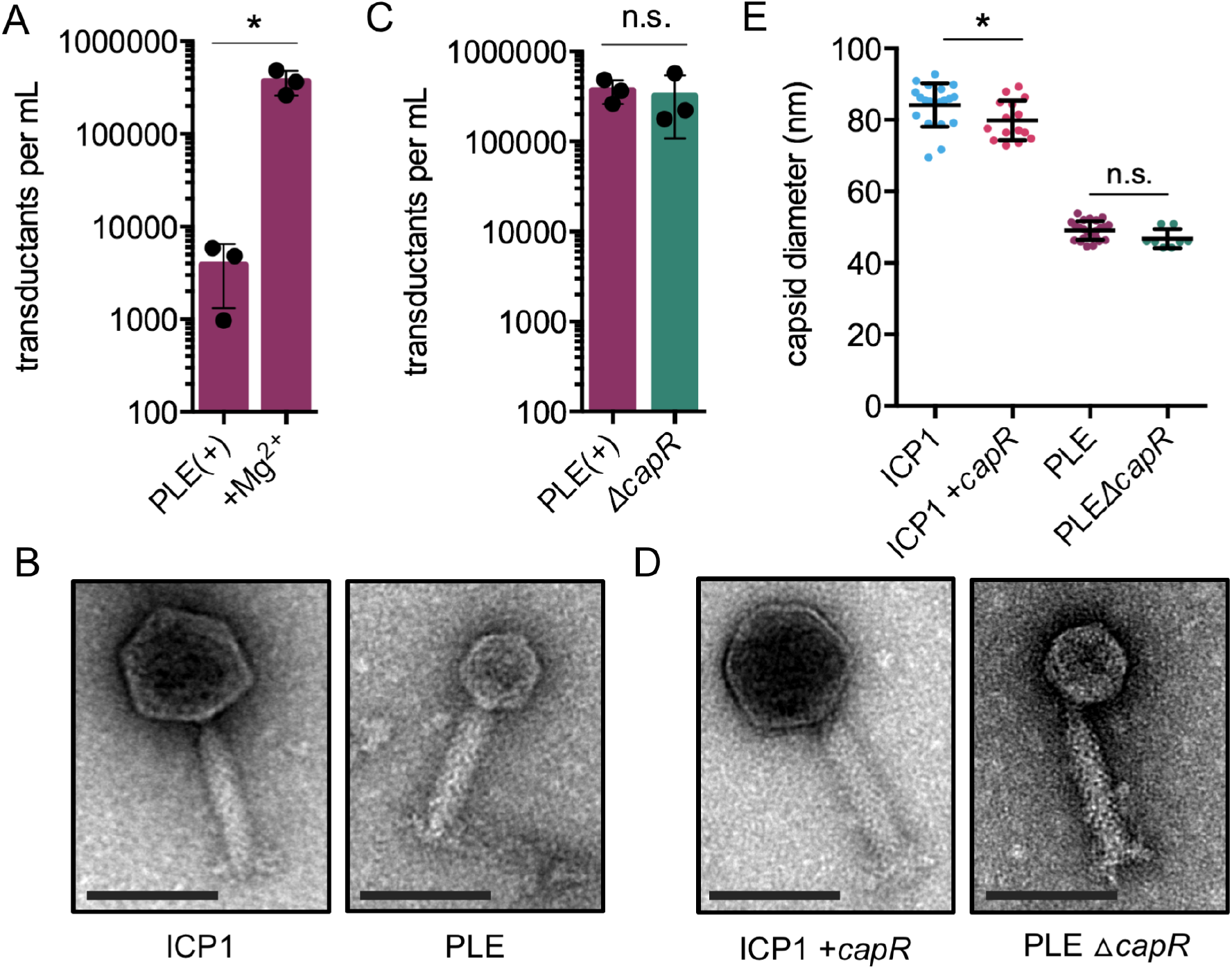
Visualization of PLE transducing particles contextualizes CapR activity. (A) Wild type PLE transduction with and without supplemental MgC1_2_, measured by transfer of a PLE marked with a kanamycin resistance gene to a PLE(−) recipient. *p <0.03 by Student t-test. (n=3) **(B)** Representative transmission electron microscopy (TEM) images of an ICPl particle and a PLE transducing particle. Scale bars = 100nm (C) Transductants quantified from PLE transduction assays described in (A) comparing wild-type PLE and PLE with the capR open reading frame deleted (PLE l::,.capR). n.s. indicates p > 0.05 by Student t-test (n=3). **(D)** Representative TEM images of an ICPl particle produced from a *V.* cholerae host strain expressing capR from the lacZ locus during infection (P,ac-capR) and a PLE transducing particle produced from PLE with the capR open reading frame deleted (PLE l::,.capR). Scale bars = 100nm. **(E)** Measurements of ICPl and PLE particle capsid diameters (nm) from particles prepared in parallel. *p=0.03, n.s. indicates p>0.05 by Student t-tests (unpaired). (n 2::8)

We were next interested to determine if CapR-mediated repression of ICP1’s capsid operon was a specific mechanism to ensure robust PLE transduction. Smaller heads would likely require fewer capsid monomers than full-sized ICP1 heads, so in the context of PLE’s life cycle the operon-specific repression could be supporting PLE transduction. However, PLE lacking *capR* transduces just as efficiently as wild-type PLE, demonstrating that CapR-mediated capsid repression activity is not essential for PLE transduction (Figure 4C). PLE transducing particles produced from a *capR* knockout also have the same morphology as wild-type PLE particles by electron microscopy (Figure 4D). Even though *capR* was not necessary for the formation of PLE particles with small heads, we unexpectedly observed that ICP1 particles produced from *V. cholerae* hosts expressing *capR* have a slightly smaller average capsid diameter (~4nm smaller than ICP1 from a wild-type host infection) (Figure 4E). This small yet significant difference could be the result of ICP1 particle instability due to CapR repression of the entire capsid operon, which also encodes other genes putatively essential for capsid morphogenesis (e.g. capsid decoration protein and scaffold). Repression of all of these components could impact ICP1 assembly, and more unstable or improperly assembled particles may experience differential staining during preparation for visualization. More quantitative microscopy methods would be necessary to determine the true magnitude and source of the observed difference. Taken together, these results suggest that capsid repression alone is not sufficient to alter ICP1’s particle head size. Although this was somewhat unsurprising given that *capR* expression in the absence of PLE has minimal impact on ICP1, it opens more questions about the precise role of capsid repression in the context of total PLE activity.

### CRISPRi knock-down of capsid operon impedes ICP1 progeny production

We were surprised to discover the limited ability of CapR to negatively impact the ICP1 lifecycle. We hypothesized that CapR-mediated repression of ICP1’s capsid operon may be tuned to this specific level to benefit PLE. PLE likely hijacks capsid morphogenesis proteins from ICP1, so while more potent or complete repression could substantially inhibit ICP1, it could also be detrimental to PLE. In order to test this, we took advantage of a native Type I-F CRISPR-Cas system in ICP1 (30) and designed a CRISPR interference (CRISPRi) system to knock down expression of the capsid operon during ICP1 infection completely independent of PLE. Briefly, a plasmid in *V. cholerae* is pre-induced to express a pre-CRISPR RNA (pre-crRNA) targeting a PAM-adjacent sequence near or within the promoter region of the operon of interest. A non-targeting spacer is expressed in a separate strain as a control. Each strain is infected with a CRISPR-Cas(+) ICP1 phage with a genetic modification to *cas2-3* that inactivates Cas2-3 nuclease activity (*cas2-3\**). Upon infection, the ICP1-encoded Csy complex is expressed and then processes and associates with the pre-expressed (now mature) crRNA. This complex binds the cognate DNA target site. Cas2-3* is recruited to the site but cannot cut the DNA, and the Csy complex blocks transcriptional machinery from accessing the promoter region. (Figure 5A).

**Figure 5.**
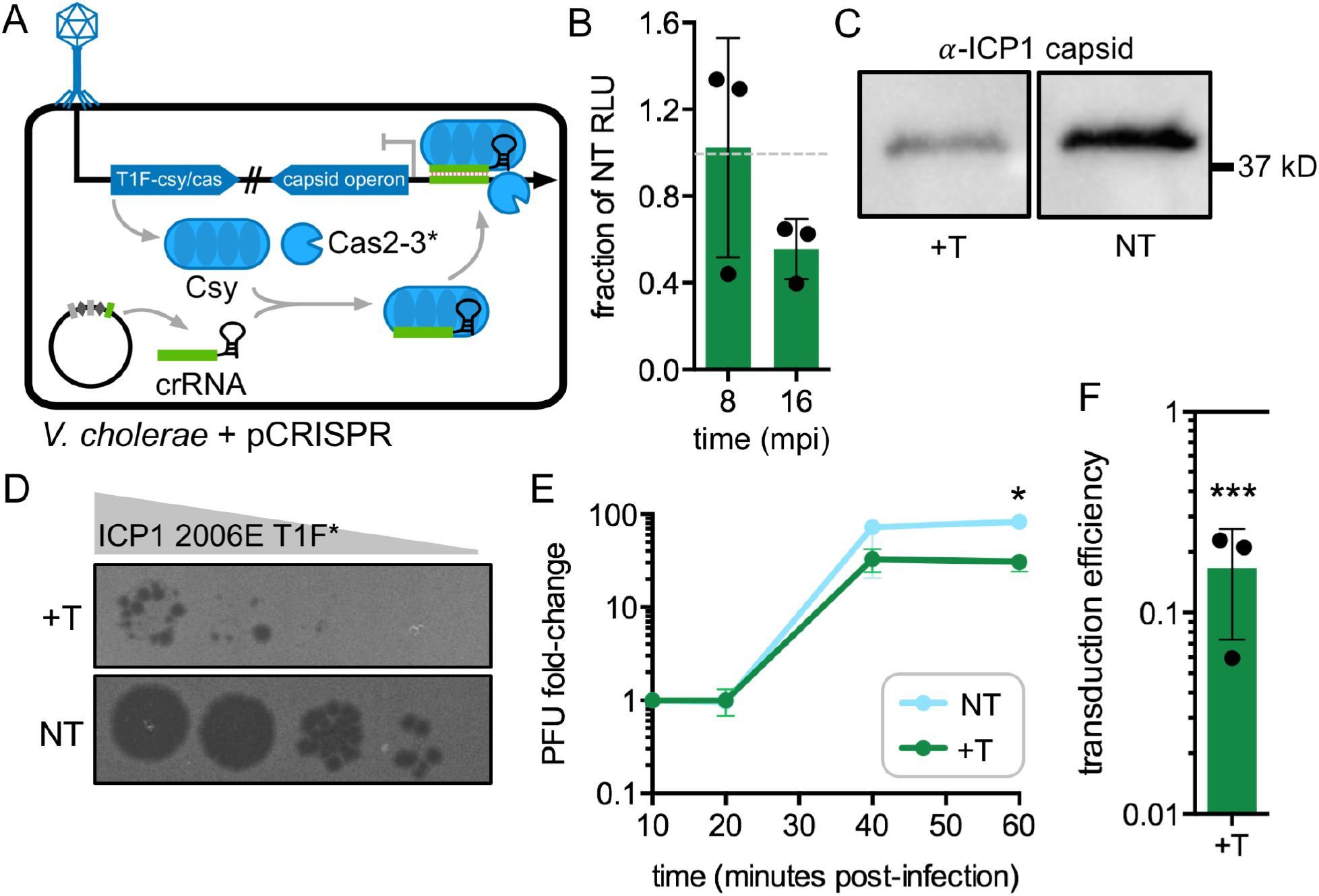
Knockdown of capsid operon by CRISPRi efficiently restricts ICPl replication. (A) Schematic of novel phage-directed CRISPRi knockdown system. ICPl expresses its native TypelF CRISPR system (TlF-csy/cas) with a nuclease-dead Cas2-3 derivative (Cas2-3*). This Csy interference complex binds a CRISPR RNA (crRNA) expressed as a pre-CRNA in the *V. cholerae* host prior to infection. The complete interference complex binds the target site of interest, blocking transcription in a sequence-directed manner. **(B)** ICPl capsid operon reporter luminescence during capsid operon CRISPRi (+T, targeting) expressed as a fraction of luminescence from a non-targeting (NT) control infection. Grey dashed line indicates expression level equivalent to non-targeting infection (relative luminescence=l). **(C)** Western Blot against ICPl capsid at 16 minutes post-infection with capsid operon CRISPRi targeting or non-targeting spacers. Total protein from cell lysate was calculated and normalized to standardize total protein input into each lane. **(D)** ICPl spot assay. ICPl-TlF* plaques (black) on *V. cholerae* host lawns (grey) expressing a spacer targeting the ICPl capsid operon promoter or a non-targeting control. Spots represent~ 10-fold serial dilutions of ICPl. **(E)** ICPl one-step burst assay quantifying the number of ICPl plaque forming units (PFU) produced over a single round of ICPl infection in a host with capsid operon CRISPRi targeting vs. a non-targeting spacer. *p=0.008 by Student t-test (n=3) **(F)** PLE transduction under capsid operon CRISPRi targeting, expressed as a fraction of transduction from a host expressing a non-targeting guide. ***p=0.0001 by Student t-test (n=3)

We applied this knockdown strategy to the ICP1 capsid operon using a crRNA targeting the −35 region of the capsid operon promoter, which was informed by transcriptomics (21). Multiple spacers within the promoter region were tested, however we only observed potent inhibition of the ICP1 capsid operon with a spacer targeting the −35 of the operon’s promoter on the template strand. Transcriptional repression under CRISPRi targeting was assessed by measuring luminescent output from the NanoLuc reporter integrated downstream of the capsid gene in the ICP1-*cas2-3\** background. After a single round of infection with CRISPRi targeting the capsid operon promoter, the amount of nanoluciferase signal was reduced to between half and one third of the non-targeting control (Figure 5B). By Western blot probing for ICP1 capsid, targeting reduced ICP1 capsid protein by approximately three-fold compared to a non-targeting control. (Figure 5C, Supplementary Figure S6B). CRISPRi targeting of the capsid operon is detrimental to ICP1: it reduced ICP1 efficiency of plaquing to approximately one-fifth of a non-targeting control and resulted in more heterogeneity in plaque size (Figure 5D, Supplementary Figure S6A). To further confirm reduction in ICP1 progeny production by CRISPRi, we performed one-step burst assays with targeting and observed a three-fold reduction in ICP1 particle formation in a single round of infection (Figure 5E). It was remarkable that the ICP1 lifecycle was dramatically impacted given that the measures of capsid operon repression by CRISPRi appeared surprisingly similar to the level of repression as we measured for CapR, which by contrast does not inhibit the production of ICP1 progeny. This suggests that CRISPRi targeting could be a more efficient means of repression, but the sensitivity of these molecular assays may not allow for a more detailed dissection of the differences that are occurring. All together, these results advocate for strict regulation of capsid operon expression as an essential component of the ICP1 lifecycle. This strict threshold level appears perturbed by CRISPRi capsid operon targeting but somehow undisturbed by CapR-mediated repression.

We next investigated whether efficient CRISPRi targeting at the ICP1 capsid operon would have an impact on PLE transduction. The crRNA-expressing plasmid was transformed into a PLE(+) *V. cholerae* strain and PLE transduction levels were determined. CRISPRi targeting reduced PLE transduction by nearly an order of magnitude (Figure 5F). The same crRNA that was detrimental to ICP1 reproduction also reduced PLE transmission, further supporting the hypothesis that PLE hijacks ICP1’s capsid morphogenesis proteins and the moderate degree of capsid repression achieved by CapR is evolutionarily optimal for PLE transmission.

### CapR is a DNA binding protein with specificity to the ICP1 capsid operon promoter

We hypothesized that CapR could act as a traditional transcriptional regulator by interacting with ICP1 DNA at the capsid operon promoter region to cause targeted transcriptional interference, but the CapR protein sequence does not contain any predicted DNA binding motif or fold. In order to decipher if the mechanism of CapR-mediated capsid operon repression involves direct DNA recognition and binding at that region, we expressed and purified CapR protein from *E. coli* and evaluated DNA binding activity *in vitro* by electrophoretic mobility shift assays (EMSAs). We first tested the ability of CapR to bind ICP1 DNA by exposing the protein to two DNA probes: probe A, which contains the entire intergenic sequence upstream of the ICP1 capsid operon (including the promoter), and probe B, which is a similar length and GC content from within *gp183-gp184* in ICP1 (Figure 6A). We observed CapR binding to probe A (containing the capsid operon promoter), but no detectable binding to probe B (Figure 6A). This confirms the DNA binding activity of CapR protein and demonstrates its specificity for the ICP1 capsid operon promoter region.

**Figure 6.**
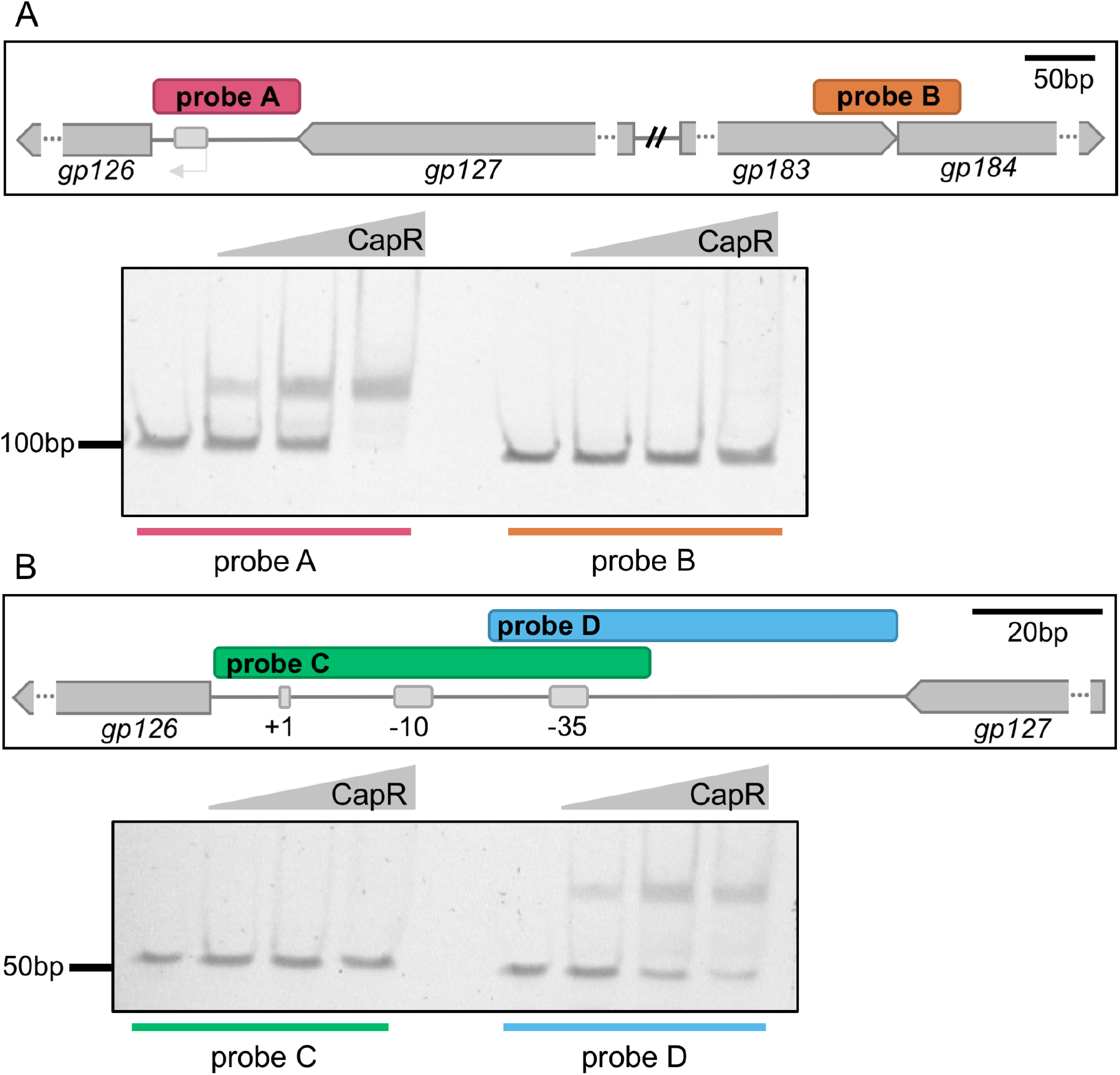
CapR is a DNA binding protein that recognizes the ICPl capsid promoter region. **(A)** Top - schematic of ICPl genomic sequence represented in probe A (the entire intergenic space between *gp127* and *gp126*, the start of the ICPl capsid operon) and probe B (a probe of similar genomic content in a distant position spanning ICPl *gpl83-gp184).* The promoter region is indicated with a light grey box and arrow. Ellipses (…) interrupting genes indicate that the gene length has been truncated for simplicity in the diagram. Bottom - electrophoretic mobility shift assays (EMSAs) following DNA binding reactions with CapR protein and probes A and B. The first lane in each probe set does not contain protein, the following lanes contain increasing concentrations of CapR protein. (B) Top - schematic of probes C and D, which each constitute half of probe A. Grey bars in intergenic region indicate capsid operon promoter elements. Scale bar represents 20bp. Bottom - EMSAs following DNA binding reactions with CapR protein and probes C and D. The first lane in each probe set does not contain protein, the following lanes contain increasing concentrations of CapR protein.

In order to further dissect the mechanism of CapR-mediated transcriptional inhibition, we determined a more precise binding site for CapR within the capsid operon promoter region by repeating CapR-DNA binding assays with a different pair of probes: probe C, which spans half of the intergenic space and includes the −10 and −35 of the promoter for the capsid operon, and probe D, which includes the −35 and extends through the other half of the intergenic space adjacent to *gp127* (Figure 6B). We detected CapR binding specifically to probe D and not probe C (Figure 6B), which indicates that CapR does not repress transcription by occluding the promoter directly. However, binding slightly upstream of the promoter region could occlude other enhancer DNA sequences or interfere with binding of transcriptional activators, which frequently bind sites upstream of the −35 (31). CapR binding upstream of the promoter could also cause additional tension during helix-unwinding that could prevent transcriptional machinery from accessing the promoter region. Overall, the data demonstrate that CapR directly binds DNA to interfere with ICP1 transcription of the capsid operon.

### CapR is related to a domain of a family of homing endonucleases

The evolutionary origin of PLE remains a mystery, but like most satellite systems, it is not closely related to its helper phage ICP1. We have characterized the PLE-encoded replication initiation factor RepA, which appears to most closely resemble proteins from Gram positive bacteria (17), while other PLE proteins like the lysis inhibition disruptor, LidI, have no apparent homologs to anything previously sequenced (18). CapR, like the majority of PLE-encoded proteins, has no recognizable bioinformatic signatures. Interestingly, this small 15.5kDa protein most closely resembles N-terminal domains of proteins from ICP1, other vibriophages, and other more distantly-related phages (Supplementary Figure S7A, Supplementary Table S1). Closer examination of the ICP1 CapR-domain-containing proteins revealed that these ICP1-encoded proteins are fairly similar to one another along their total lengths. They all also contain a conserved C-terminal domain annotated as “T5orf172” in Uniprot or a domain with appreciable identity to T5orf172 domains (Figure 7A). This particular domain has been subjected to little exploration, but it has been identified as often co-occurring with the conserved N-terminal DNA-binding domains of Bro and KilA proteins (7). Bro-N domains in particular are so prevalent in the genomes of DNA viruses that they have been classified as their own DNA-binding superfamily. Therefore, by a combination of “guilt by association” and homology to these more well-classified domains, T5orf172 domains are also predicted to have DNA binding activity.

**Figure 7.**
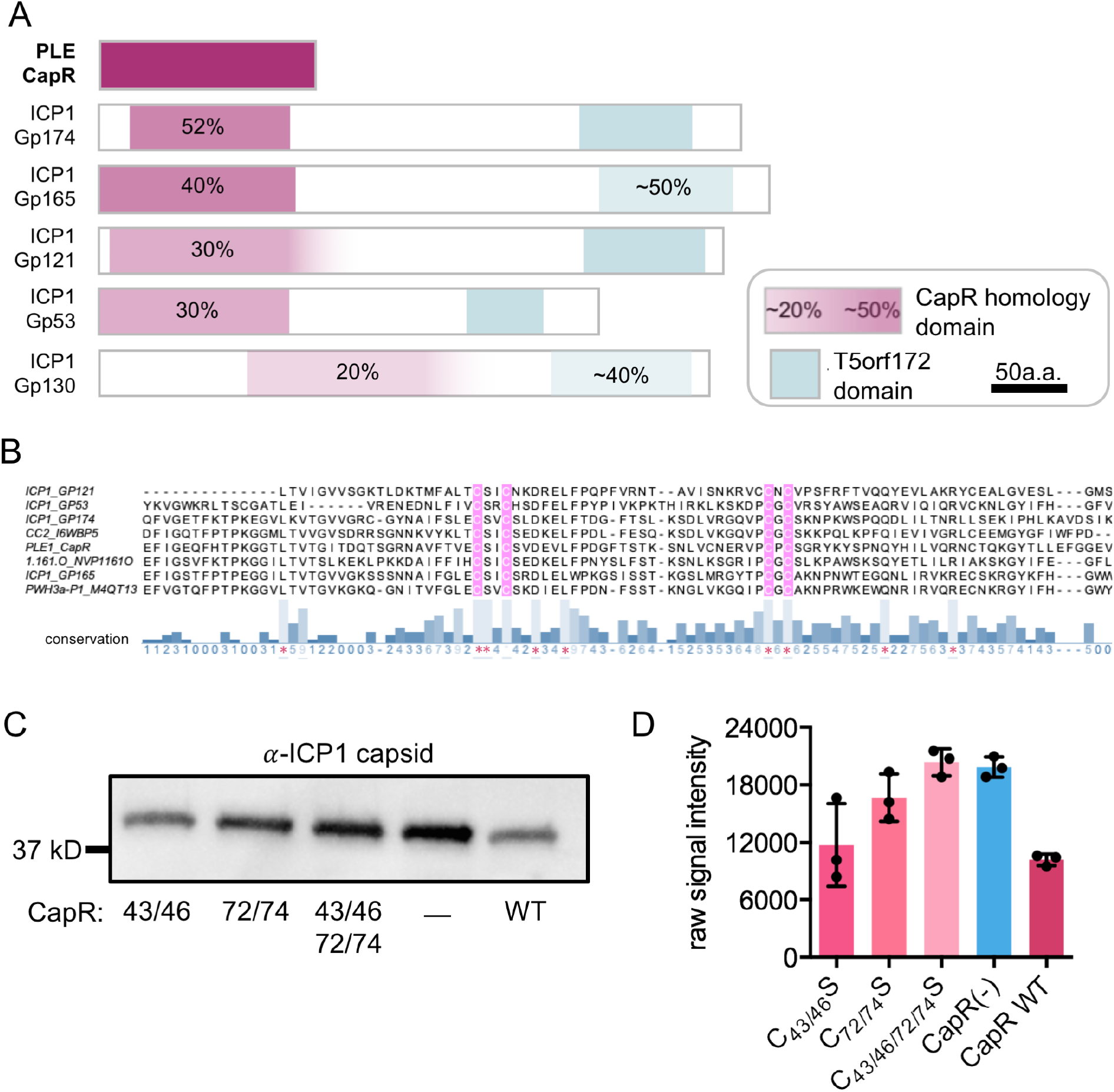
CapR is related to a lineage of homing endonucleases. **(A)** Visual representation of CapR protein homology domains in ICP1_2006_E gene products(Gp) predicted to be homing endonucleases. Purple segment denotes portion of the gene that aligns with CapR, faded shading indicates that the alignment persists but degenerates in quality/homology along the homologous portion. **(B)** Trimmed multiple sequence alignment of N-terminal domains with highest(30-50%) homology to CapR. Pink highlighting denotes conserved cysteine reside sets. Conservation score is noted underneath with blue bars. **(C)** Western blot against ICPl capsid protein 16 minutes post-infection in *V cholerae* hosts expressing the wild-type CapR allele(WT), or alleles with sets of conserved cysteines mutated to serines(numbers denote C-S substitution positions), or an empty vector control(−). **(D)** Quantification of triplicate Western blot experiments represented in(C).

A deeper examination of the domains associated with T5orf172, Bro-N, and KilA-N revealed additional insights into the particular origins of CapR and related ICP1-encoded proteins. Bro-N and KilA-N DNA-binding domains are often found in proteins that also contain GIY-YIG endonuclease domains. The T5orf172 domain itself is also categorized in the same superfamily as GIY-YIG domains. GIY-YIG domains are found within the catalytic site of a subfamily of homing endonucleases (HEGs), mobile genetic elements that encode nucleases that cut at a target site to facilitate the insertion of a copy of the HEG gene at that target site (11). A BLASTp search for proteins with identity to the ICP1 proteins with CapR homology domains (in ICP1_2006_E: Gp174, Gp165, Gp121, Gp53, Gp130) revealed several similar predicted HEGs in other viruses. In particular, both Gp165 and Gp130 have similarity to a putative endonuclease Orf141 in *E. coli* phage T5 (Supplementary Figure S7Bi, and Bii). Gp130 also contains portions similar to HEGs in *Bacillus*-phages AR9 and PBS1 and Actinophage K2 (Supplementary Figure S7Bi). Gp121 resembles putative endonucleases from Lausannevirus and *Escherichia-*phage, as well as a *Bacillus*-phage group I intron protein (a self-splicing HEG-relative) (Supplementary Figure S7Biii) Although none of the ICP1 genes have the canonical endonucleolytic motif, these bioinformatic signatures, abundance of similar genes, and flexibility of motifs composing the GIY-YIG active site led us to hypothesize that this collection of genes could represent ICP1 HEGs.

CapR shares varying degrees of homology with the N-terminal region of these putative HEGs, so we next attempted to identify residues critical to CapR function by looking for conservation among the corresponding N-terminal domains of the phage HEGs. In a trimmed multiple sequence alignment, we identified two sets of conserved cysteine residue pairs spaced 24-26 amino acids apart (CxxC ← 24-26 → CxC) within the domain (Figure 7B). These residues are predicted in some sequences to be nested between beta sheets (Supplementary Figure S8A). This arrangement is reminiscent of a zinc finger in the DNA binding domain of the homing endonuclease iTev-I (a distant relation, but the most well-characterized protein somewhat similar to ICP1 putative HEGs), which contains a small degenerate treble-clef fold with four similarly-arranged cysteines coordinating a zinc ion. The zinc finger subdomain makes two direct hydrogen bond contacts with DNA, suggesting that the subdomain may be capable of independent DNA binding and that zinc coordination by the four cysteine residues is likely critical to that activity (32).

To test if these conserved sets of cysteine residues coordinate CapR-DNA binding, each set of cysteines in CapR was mutated to serines, as well as a mutant with all four residues mutated, for a total of three mutant strains: C_43/46_S, C_72/74_S, and C_43/46/72/74_S. Mutant CapR was expressed during ICP1 infection and capsid protein production was assessed by Western blot (Figure 7C and D). With each successive CapR mutation we observed decreased repression of ICP1 capsid protein. The first set of cysteines appear to be less involved in coordination as the C_43/46_S mutant displays only a slight relief in capsid repression compared to the wild-type allele, while the C_72/74_S mutant allowed more ICP1 capsid expression. Infection of the C_43/46/72/74_S mutant resulted in no detectable capsid repression (Figure 7C and D), indicating that this mutant allele can no longer mediate repression at the capsid operon. Each of the mutant *capR* alleles appeared to retain some degree of capsid repression activity as measured by NanoLuc reporter assay, however none repressed as efficiently as wild-type CapR, with the C_43/46/72/74_S mutant showing the greatest degree of repression relief (Supplementary Figure S9C). Taken together, these results validate that the four highly conserved cysteine residues in CapR are crucial for its DNA binding and are likely forming a zinc finger fold to aid in this activity.

## Discussion

Here, we explore the mechanism and implications of PLE-mediated targeted repression of the ICP1 capsid operon. CapR, a conserved PLE gene product with no predicted function, is both necessary and sufficient to achieve the complete extent of ICP1 capsid operon repression observed with wild-type PLE. CapR achieves transcriptional repression activity by direct DNA binding within the intergenic space containing the capsid operon promoter region. CapR’s DNA binding activity was not supported by direct sequence similarity to known DNA binding motifs, but the protein has similarity to domains of proteins in ICP1. These ICP1 proteins are putative homing endonucleases, which are proteins that contain multiple domains that all directly interact with DNA (33). A putative zinc finger motif conserved among CapR and putative ICP1 homing endonucleases containing CapR-like sequence appears to contribute to the protein’s DNA binding activity.

The strategy of limiting phage late gene expression appears to be a common feature of phage satellite biology, though the implications of such restriction beyond general phage interference remain largely unknown. P4, a phage satellite in *E. coli*, encodes a mimic (Delta) of its helper phage P2’s late gene transcriptional activator (Ogr) in order to modulate transcription by direct competition with the phage regulator (34). Interestingly, Ogr homologs contain sets of conserved cysteines indicative of a zinc finger structure similar to CapR. In some SaPI satellites, transcriptional repression of late helper phage genes is achieved by expression of a very potent repressor, PtiA, whose activity must be reduced by a co-expressed protein, PtiM, in order to allow enough structural gene expression for SaPI transmission (20,35). In this system, satellite-mediated repression of phage late genes is thought to serve as a direct phage restriction mechanism. By contrast, PLE achieves a very similar transcriptional degree of repression using a single gene product. The incomplete transcriptional repression by CapR dispenses with the need for other regulatory components for this particular phenomenon, distinguishing its mechanism from the transcriptional repression employed by SaPIs. Another factor unique to PLE’s CapR is its lack of ability to inhibit the phage lifecycle, indicating that its purpose in total PLE activity is distinct from phage interference.

We were surprised to discover that CapR-directed repression of the ICP1 capsid operon alone did not impair ICP1 progeny production. However, the repression imposed by CapR is not complete; there is an approximately three-fold reduction in transcriptional activity from the operon resulting in roughly one third as many capsid morphogenesis proteins available for ICP1 to construct progeny capsids. This suggests that ICP1 capsid operon components are synthesized in great excess during a single round of ICP1 infection and that fewer than half of these proteins are incorporated into progeny phage heads. Excessive structural gene synthesis could be a mechanism employed by ICP1 to ensure complete host take-over by saturating host protein synthesis machinery with phage component transcription and translation. This is supported by the observation that the capsid operon is the most highly-expressed in the ICP1 transcriptome late in infection (21), reflecting a general pattern of excess structural protein production in cytotoxic viruses (36). If only a subset of structural components is destined to be assembled into complete phage particles, this could explain ICP1’s lack of sensitivity to capsid repression alone outside the context of total PLE activity. ICP1 infections were conducted in nutrient-replete conditions, so perhaps CapR-mediated repression could have negative impacts in an infection context with different phage kinetics or nutrient availability. When the ICP1 capsid operon was transcriptionally repressed by an alternative CRISPRi strategy, the repression was detrimental enough to ICP1 to reduce efficiency of plaquing and burst size. Curiously, our measurements of capsid production by Nanoluc reporter and Western blot appear roughly equivalent between CRISPRi and CapR-mediated operon repression. This could be because the degree of repression exerted by CapR is already at the limit of detectable difference in this biological context. Examining an alternative knockdown strategy enriched our understanding of essential phage genes but opened questions about why PLE would not favor stronger CapR-mediated repression over evolutionary time if PLE is already employing other ICP1 inhibition strategies.

CapR acts as an inefficient repressor, but this perceived shortcoming is perhaps its most useful feature. Its role in the context of PLE exemplifies the delicate balance that phage satellites must achieve in order to transfer horizontally: the helper phage lifecycle must be sufficiently limited in order to favor satellite propagation, but certain helper phage genes and/or processes are required for satellite propagation, including packaging of the satellite genome into helper phage structural components. We show that repression of the phage capsid operon by CapR allows for complete PLE transduction, while a more restrictive mode of repression (CRISPRi) also restricts PLE transduction. This exemplifies the delicate threshold of repression achieved by CapR, allowing it to perform transcriptional interference without impeding PLE transmission.

The first visualization of PLE transducing particles reported here revealed that PLE mobilizes in ICP1-like particles with smaller heads likely requiring fewer capsid subunits. Similar to the general strategy of repressing inducing phage late gene expression, the majority of characterized viral satellites modify their virions by reducing capsid head size (13). This places PLE and ICP1 squarely within the paradigm of phage-satellite pairs with reduced head diameter morphology and explains how PLE can utilize these components while simultaneously repressing their expression via CapR. Particle size redirection is an efficient phage interference mechanism, as phage genomes are too large to be completely packaged in modified satellite particles. However, because capsids generally require other structural components to direct assembly into the proper shape and size, reducing the available capsid monomer pool is not sufficient to favor formation of smaller particles. SaPIs and P4 both encode alternative scaffolds that force capsid monomers into configurations with smaller diameter and volume. Future work will investigate the PLE-mediated mechanism of particle head size remodeling.

We previously demonstrated that all five known PLEs are able to repress ICP1 capsid transcription and translation to similar degrees (21). This observation coupled with the high degree of conservation between CapR alleles in these other PLEs (Supplementary Figure S8B) imply a highly conserved role for CapR in capsid repression among all PLEs. Because there is slight variance between these alleles and their protein product’s activity could feasibly be escaped with genetic mutation, we examined whether any sequenced ICP1 isolates displayed evidence of such selection at the predicted CapR binding site upstream of the ICP1 capsid operon. We were surprised to see that all previously sequenced ICP1 isolates contain 100% identical intergenic regions upstream of the ICP1 capsid operon, suggesting that the variations in CapR alleles from different PLEs are not likely “responding” to changes in the recognition sequence, but may instead be variations with different degrees of stability or simply the consequence of genetic drift. It is also interesting that the ICP1 capsid operon promoter region is highly conserved. This could be an indication that whatever native ICP1 protein is responsible for late gene transcriptional regulation is also highly conserved and reliant on the fidelity of its recognition site.

The similarity of CapR to putative ICP1 homing endonucleases brings up an intriguing hypothesis about the potential evolutionary origin of a *capR-*like allele in an ancestral PLE. Similar to P4’s Delta gene product which directly competes with helper phage P2’s activator Ogr for binding, CapR could possibly accomplish transcriptional repression by mimicking a native ICP1 transcriptional activator. However, little is currently known about transcriptional control within the ICP1 genome, and ICP1 proteins that contain sequence most similar to CapR (Gp174 and Gp165) are likely HEGs. Gp174 and Gp165 could theoretically be active HEGs, but this would require that most ICP1 genomes lack their target cut sites. It is possible they have accumulated mutations resulting in loss of endonuclease activity. The ICP1 HEGs could also have been neo-functionalized to serve roles in ICP1 processes such as transcriptional regulation. The HEG iTev-I in T4 serves a dual regulatory role as an autorepressor by recognizing an alternative binding site within the T4 late promoter driving its expression, performing genetic regulation independent of nuclease activity (37). An active homing endonuclease in the ICP1 genome could perform a similar moonlighting activity. The HEG could home within the genome and acquire mutations, either to escape HEG targeting or degenerate over rounds of replication. Some of these mutations could impact nuclease target recognition, allowing for the HEG pseudo-copy to home into a new location in the phage genome. Because these genes are only expressed in a *V. cholerae* host during phage infection, there is opportunity for an ancestral PLE to encounter and survive targeting by one of the diversified ICP1 HEGs. A HEG hosted by PLE could acquire mutations and even lose entire domains until a neo-functionalized version of the allele emerged. These HEGs present a type of evolutionary shortcut for PLE, as they are likely already adapted to interacting with the ICP1 genome. The modern *capR* allele could then be easily deployed against the very genome it potentially originated from. This hypothetical chain of events illustrates the potential rich evolutionary history of a single conserved PLE gene that was likely acquired in an ancestral PLE.

This work provides the first example of putative HEG-mediated transcriptional network evolution. A handful of related discoveries in other systems support the role of MGEs as critical modulators of transcriptional control in the context of evolution. In the mammalian placenta, integration of endogenous retroviruses around regulatory regions drives rapid evolution of transcriptional networks (8). In a more closely-related bacterial system, *B. subtilis* encodes Rok, a transcriptional regulator involved in repressing competence gene expression and regulating other unrelated bacterial functions (38). Rok is also responsible for silencing a specific integrated MGE and is found bound to large foreign A/T-rich DNA regions common to MGEs, indicating a tight evolutionary constraint placed on bacterial gene content and control by a mechanism likely intended to restrict MGEs. Rok’s involvement in the regulation of bacterial genes that did not likely originate from MGEs indicates that some of those genes could have originally been delivered into the host by MGEs and neo-functionalized by the host. CapR’s putative neo-functionalization from an active homing endonuclease in the ICP1 genome was likely expedited by providing a fitness advantage to any allele that was no longer able to target and cut the PLE or *V. cholerae* host genome, but perhaps still maintained some activity against their common enemy: ICP1. Investigation of CapR’s origin opens many other questions about the MGE-facilitated evolution of similar transcriptional cross-talk. Phage satellites in particular are likely to contain a wealth of such undiscovered phage interference mechanisms that evolve in novel ways as they stand boldly in the midst of bacteria-phage arms races.

## Data Availability

Uniprot IDs for all proteins in text are listed in Supplementary Table S1.

Transcriptomics data (21)are deposited in the Sequence Read Archive under the BioProject accession [PRJNA609114]

ICP1_2006E Genome GenBank: MH310934.1

## Acknowledgements

We would like to thank all current and former members of the Seed Lab for contributing helpful ideas and feedback, particularly Steph Hays for graphic and content advisement, James Mao for initial EMSA troubleshooting, Zach Barth for useful discussions and editing, Kristen LeGault for the initial discovery of particle stabilization by magnesium as well as useful discussions and editing, and Maria Nguyen for strain construction. Special thanks to the staff at the University of California Berkeley Electron Microscope Laboratory for advice and assistance in electron microscopy sample preparation and data collection. This work used the Vincent J. Proteomics/Mass Spectrometry Laboratory at UC Berkeley, supported in part by National Institutes of Health S10 Instrumentation Grant [S10RR025622].

