## Supplemental tables and figures for "A phage satellite tunes inducing phage gene expression using a domesticated endonuclease to balance inhibition and virion hijacking"

**Supplementary Table 1. Proteins listed in figures/text**

| Name in text | Uniprot ID | Annotation | Source organism |
|---|---|---|---|
| ICP1 Gp174 | A0A385EI32 | T5orf172domain containing protein | ICP1_2006E |
| ICP1 Gp165 | F1D1I8 | uncharacterized protein | ICP1_2006E |
| ICP1 Gp53 | F1D175 | T5orf172domain containing protein | ICP1_2006E |
| ICP1 Gp121 | F1D1E4 | T5orf172domain containing protein | ICP1_2006E |
| ICP1 Gp130 | F1D1F3 | uncharacterized protein | ICP1_2006E |
| PWH3a-P1 | M4QT13 | T5orf172 domain-containing protein | Vibrio phage PWH3a-P1 |
| 1.161.O | A0A2I7RCZ8 | uncharacterized protein | Vibrio phage 1.161.O._10N.261.48.C5 |
| 1.170.O | A0A2I7REZ2 | Meiotically up-regulated protein 113 | Vibrio phage 1.170.O._10N.261.52.C3 |
| 1.193.O | A0A2I7RKM4 | uncharacterized protein | Vibrio phage 1.193.O._10N.286.52.C6 |
| S4-7 | A0A1C9LW33 | T5orf172 domain-containing protein | Vibrio phage S4-7 |
| CC2 | I6WBP5 | uncharacterized protein | Aeromonas phage CC2 |
| p ¼ | A0A088C4G7 | uncharacterized protein | Shewanella sp. phage 1/4 |
| 11895-B1 | M4QUS6 | T5orf172 domain-containing protein | Vibrio phage 11895-B1 |
| T5.141 put. endonuclease | Q6QGE6 | Putative endonuclease | E. coli phage T5 |
| ExoABC | G4KK90 | Exonuclease ABC C subunit | Yersinia phage phiR1-37 |
| PBS1 HEG | A0A223LD54 | Homing endonuclease | Bacillus phage PBS1 |
| K2 HEG | A0A3S6PTV0 | Putative Hef-like homing endonuclease | Kelbsiella phage 1611E-K2-1 |
| AR9 HEG | A0A172JI54 | Hef-like homing endonuclease | Bacillus phage AR9 |
| Bobb Group I intron | A0A076G7J2 | Group I intron protein | Bacillus phage Bobb |
| LAU put. RE | F2WL88 | Putative restriction endonuclease | Lausannevirus |
| vB put. endonuclease | A0A499PY34 | Putative endonuclease | Escherichia phage vB_EcoM-Ro121c4YLVW |

**Supplementary Table 2. Strains used in this study**

| Name in text | Strain | Description | Source |
|---|---|---|---|
| PLE(-) (Figures 1, S2, S3A) | KDS6 | *V. cholerae* E7946: O1 El Tor, streptomycin resistant, CTX(+), | Lab collection |
| PLE(+) (Figures 1, S2, S3A) | KDS36 | *V. cholerae* E7946, PLE1 integrated between VCA0329 and VCA0330 | O'Hara et. al (2017) (15) |
| PLE(-) | KDS116 | *V. cholerae* E7946 Δ*lacZ*, empty cassette under IPTG-inducible ($P_{tac}$) and theophylline-inducible riboswitch (RiboE) control and kanamycin resistance marker integrated into the *lacZ* locus | McKitterick & Seed (2018) (16) |
| PLE(+) | KDS103 | *V. cholerae* E7946, PLE1 with kanamycin resistance marker downstream last ORF in PLE | O'Hara et. al (2017) (15) |
| PLE(-) Ptac-*capR* | KDS118 | *V. cholerae* E7946, Δ*lacZ*, *capR*$^{PLE1}$ under $P_{tac}$ and RiboE inducible control and kanamycin resistance marker integrated into the *lacZ* locus | This study |
| PLE(+)Δ*capR* | KDS149 | *V. cholerae* E7946, PLE1 with kanamycin resistance marker downstream last ORF in PLE, frt scar in place of *capR* (*orf2*$^{PLE1}$) | Hays & Seed (2020) (18) |
| PLE transduction recipient | KS882 | *V. cholerae* E7946, Δ*lacZ*, spectinomycin resistance marker integrated into the *lacZ* locus | O'Hara et. al (2017) (15) |
| +T | KS1743 | *V. cholerae* E7946, pCRISPR phi (plasmid harbors a synthetic BsaI-editable ICP1 CRISPR array downstream under $P_{tac}$-inducible control on a pMMB backbone) encoding a spacer targeting the -35 region of the capsid operon promoter template strand, encodes ampicillin resistance (ampR) | This study |
| NT | KS1832 | *V. cholerae* E7946 pCRISPR phi plasmid encoding a non-targeting spacer, $P_{tac}$-inducible | This study |
| +T (transduction) | KS1836 | *V. cholerae* E7946, PLE1 with kanamycin resistance marker, pCRISPR phi (ampR) encoding a spacer targeting the -35 region of the capsid operon promoter template strand, $P_{tac}$-inducible | This study |
| NT (transduction) | KS1834 | *V. cholerae* E7946, PLE1 with kanamycin resistance marker, pCRISPR phi plasmid (ampR) encoding a non-targeting spacer, $P_{tac}$-inducible | This study |
| $C_{43/46}S$ | ZN173 | *V. cholerae* E7946, Δ*lacZ*, *capR*$^{PLE1}$ with $C_{43}S$ and $C_{46}S$ substitution mutations under $P_{tac}$ and RiboE inducible control and kanamycin resistance marker integrated into the *lacZ* locus | This study |
| $C_{72/74}S$ | ZN167 | *V. cholerae* E7946, Δ*lacZ*, *capR*$^{PLE1}$ with $C_{72}S$ and $C_{74}S$ substitution mutations under Ptac and RiboE | This study |

| | | | |
|---|---|---|---|
| | | inducible control and kanamycin resistance marker integrated into the *lacZ* locus | |
| $C_{43/46/72/74}S$ | ZN170 | *V. cholerae* E7946, Δ*lacZ*, *capR*$^{PLE1}$ with $C_{43}S$, $C_{46}S$, $C_{72}S$, and $C_{74}S$ substitution mutations under Ptac and RiboE inducible control and kanamycin resistance marker integrated into the *lacZ* locus | This study |
| ICP1 | | ICP1_2006_E ΔCRISPR ΔCas2-3 | McKitterick & Seed (2018) (16) |
| ICP1-gp68NL | | ICP1_2006_E ΔCRISPR ΔCas2-3, *nanoluciferase* gene with independent translational start inserted downstream of *gp68* | This study |
| ICP1-gp122NL | | ICP1_2006_E ΔCRISPR ΔCas2-3, *nanoluciferase* gene with independent translational start inserted downstream of *gp122* | This study |
| ICP1-T1F* | | ICP1_2006_E ΔCRISPR, Cas2-3* mutation D112A inactivates nuclease activity of Cas2-3, *nanoluciferase* gene with independent translational start inserted downstream of *gp122* | This study |

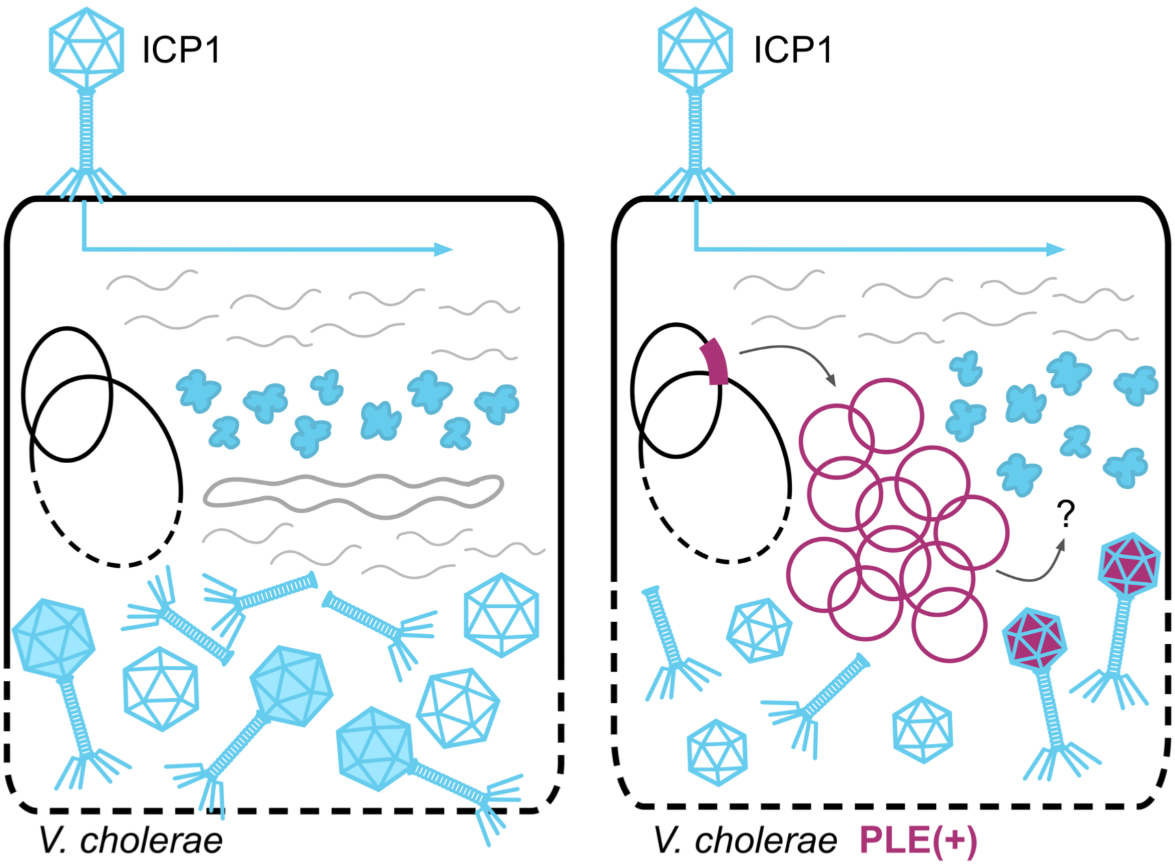


**Supplementary Figure 1. ICP1 infection of PLE(-) and PLE(+) *V. cholerae*.** Left - in a PLE(-) *V. cholerae* host, ICP1 acts as a typical lytic phage. ICP1 attaches and injects its genome. Early gene expression facilitates phage genome replication and host cell takeover. Late gene expression results in synthesis of phage structural components. The host cell chromosomes are degraded. ICP1 genomes are packaged into heads, tails are attached, and the host cell lyses, freeing progeny phage to infect neighboring cells. Right - in a PLE(+) *V. cholerae* infection, attachment, injection, and early gene expression occur as normal. PLE (purple) is triggered to excise from the chromosome and begins its own genome replication. While many PLE activities remain a molecular mystery, the sum total completely restricts formation of ICP1 progeny particles. Instead, PLE appears to re-direct ICP1 packaging material to transmit its own genome. The host cell is lysed on an accelerated timeline.

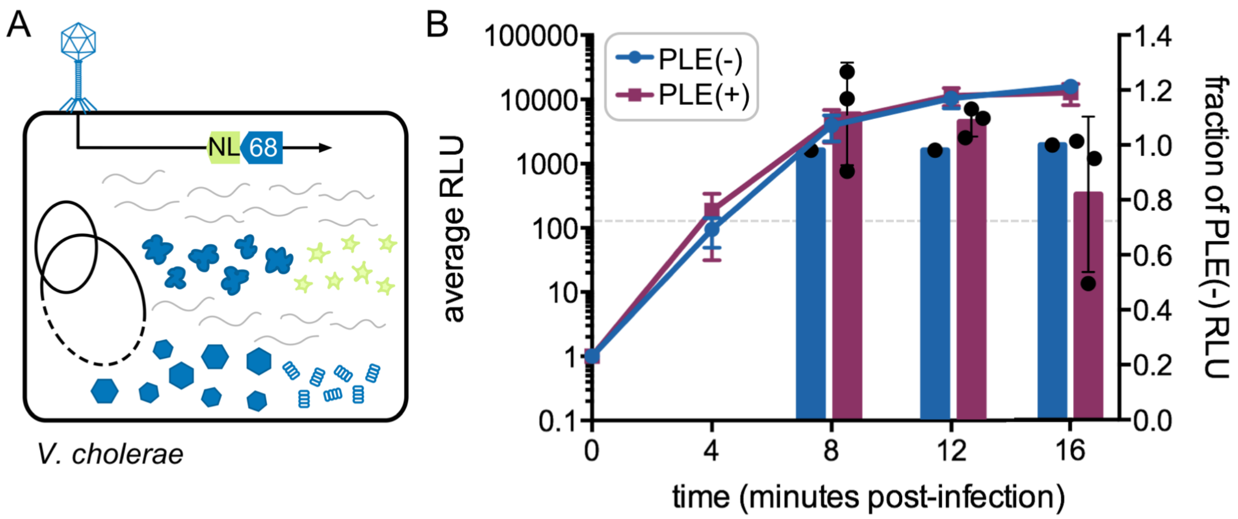


**Supplementary Figure 2. Temporal expression of ICP1 middle gene nanoluciferase reporter tool.** (**A**) Schematic of the middle (*gp68*) ICP1 gene nanoluciferase (NL) expression reporter construct. The nanoluciferase is synthesized as its own open reading frame under the native *gp68* promoter. (**B**) Left - line graph plots raw luminescent output at the timepoints indicated post-infection with the ICP1 *gp68* reporter phage in PLE(-) and PLE(+) *V. cholerae* host strains. Right - bar graphs express the raw luminescence as a fraction of the luminescent output of a PLE(-) infection. Grey dashed line indicates RLU limit of detection.

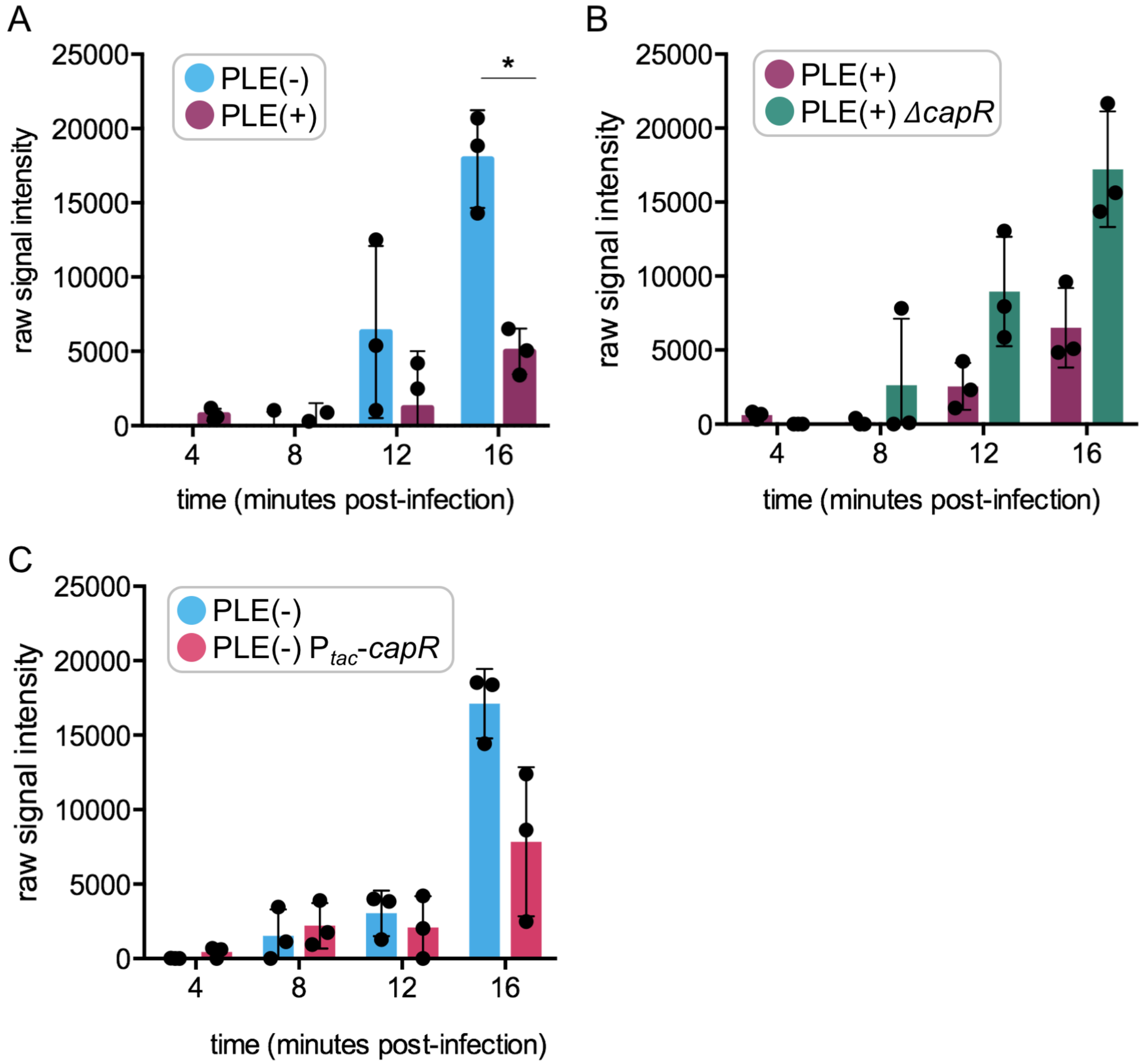


**Supplementary Figure 3. Western blot quantifications.** Signal intensity quantification (background-subtracted) for 3 biological replicate Western blot infection time-course experiments measuring ICP1 capsid monomer production in the following: (**A**) PLE(-) vs. PLE(+) *p=0.004 by Student t-test (n=3) (**B**) PLE(+) wild type vs. PLE with the *capR* open reading frame deleted (Δ*capR*) (**C**) PLE(-) vs. PLE(-) expressing *capR* from the *V. cholerae lacZ* locus (P$_{tac}$-*capR*).

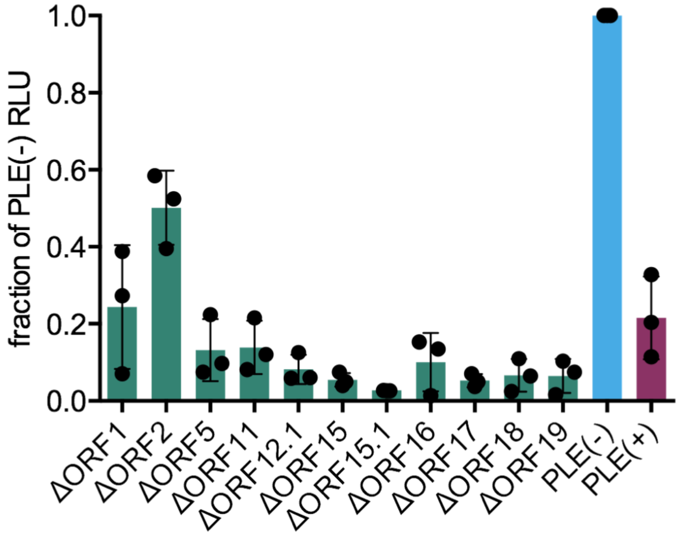


**Supplementary Figure 4. Screen of deletions of conserved core PLE genes for relieving capsid repression.** 11 PLE(+) strains, each with a single core gene knocked out, were infected with the ICP1 capsid operon reporter phage (MOI = 0.1). Luminescence output at 16 minutes post-infection is expressed as a fraction of PLE(-) reporter infection luminescence. Bars represent an average of 3 biological replicates.

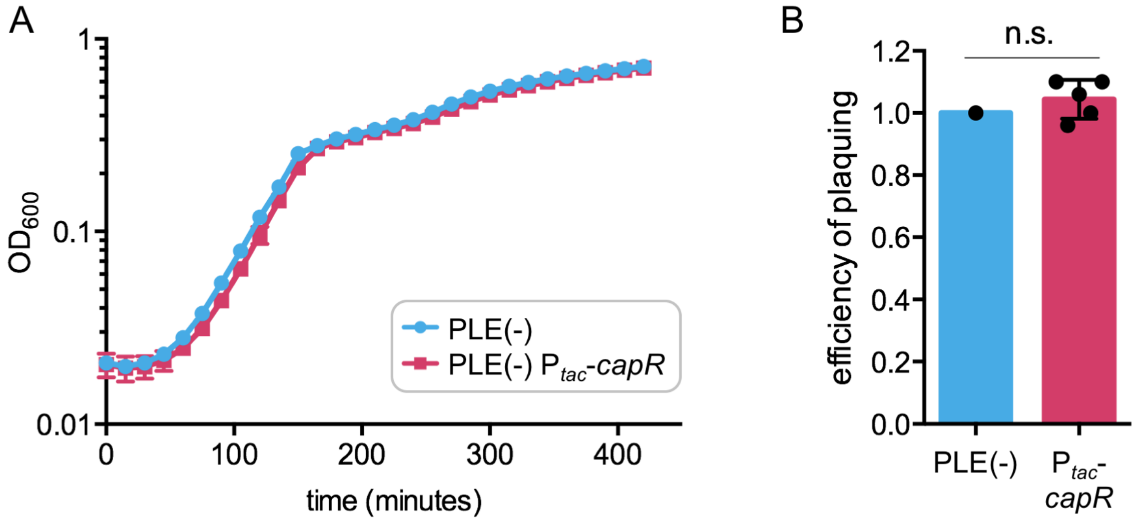


**Supplementary Figure 5. CapR expression does not interfere with *V. cholerae* growth or ICP1 efficiency of plaquing.** (**A**) *V. cholerae* (PLE(-)) growth assay with induced *capR* expression ($P_{tac}$-*capR*) or empty vector in the *V. cholerae lacZ* locus, measuring $OD_{600}$ over time. Graph represents averages from 3 biological replicate experiments conducted in technical triplicate. (**B**) Bar graphs quantify the efficiency of plaquing, or the number of ICP1 plaques from infection on the *capR*-expressing host strain ($P_{tac}$-*capR*) divided by the number of plaques from a wild type PLE(-) infection. n.s. indicates $p > 0.05$ by Student t-test. (n=5)

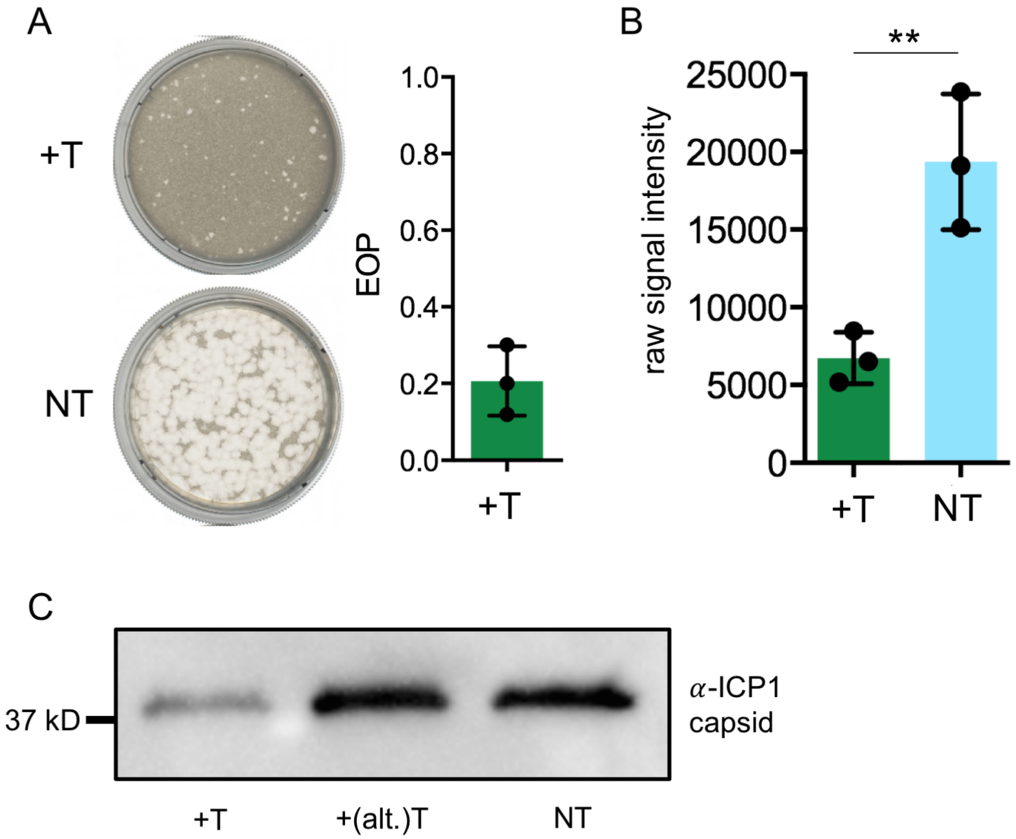


**Supplementary Figure 6. CRISPRi quantifications.** (**A**) Left - representative plaque plates at equivalent dilutions of ICP1-T1F* (clear zones) on host lawns (dark beige) expressing a spacer targeting the ICP1 capsid operon (+T) or a non-targeting spacer control (NT). Phage added at equivalent concentration on both plates. Right - bar graph enumerates the efficiency of plaquing as a ratio of ICP1-T1F* plaques on the capsid operon targeting host vs. non-targeting host. (**B**) Signal intensity quantification (background-subtracted) for 3 biological replicate Western blot infection experiments measuring ICP1 capsid monomer protein 16 minutes post-infection. **p=0.0095 by Student t-test (n=3) (**C**) Uncropped version of Western blot image in Figure 5E. +(alt)T denotes a strain expressing a different crRNA spacer that did not effectively knock down capsid expression.

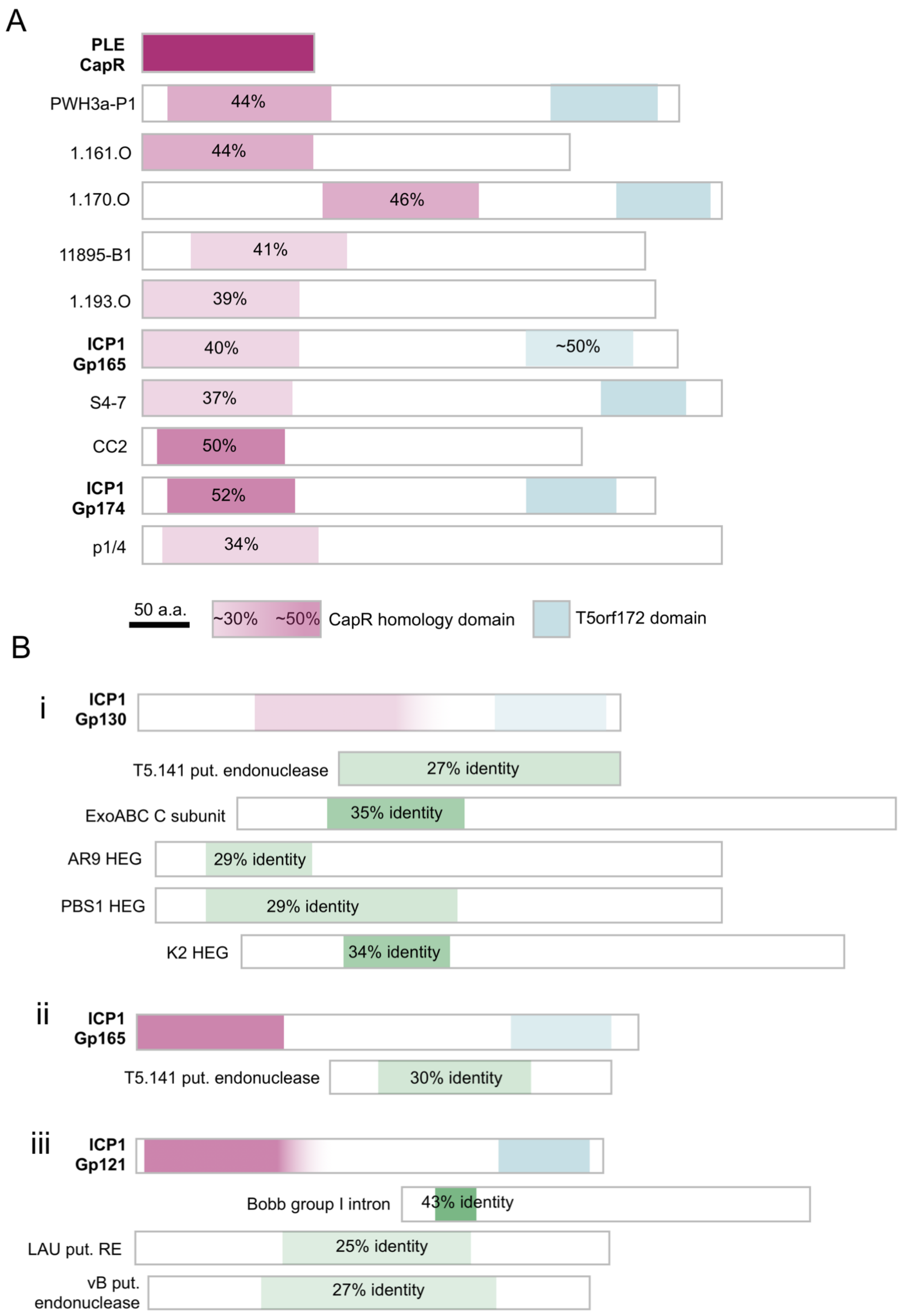
A
PLE CapR
PWH3a-P1
44%
1.161.O
44%
1.170.O
46%
11895-B1
41%
1.193.O
39%
ICP1 Gp165
40%
~50%
S4-7
37%
CC2
50%
ICP1 Gp174
52%
p1/4
34%
50 a.a.
~30%
~50%
CapR homology domain
T5orf172 domain
B
i
ICP1 Gp130
T5.141 put. endonuclease
27% identity
ExoABC C subunit
35% identity
AR9 HEG
29% identity
PBS1 HEG
29% identity
K2 HEG
34% identity
ii
ICP1 Gp165
T5.141 put. endonuclease
30% identity
iii
ICP1 Gp121
Bobb group I intron
43% identity
LAU put. RE
25% identity
vB put. endonuclease
27% identity

**Supplementary Figure 7. CapR protein homology.** (**A**) Visual representation of CapR protein homology domains among all sequenced organisms. Purple segment denotes portion of the gene that aligns with CapR and degree of shading/percentages indicate average percent identity to CapR. Protein is listed simply by host phage (except ICP1 genes), Uniprot IDs are listed in Supplementary Table S1. (**B**) Select ICP1 putative homing endonuclease genes (HEGs) and their homology to other HEGs. Green shading and bar positioning indicate the approximate position and degree of alignment to the ICP1 HEG at the top of each box (i-iii), and average percent identity is specified in the bar. LAU put. RE = putative restriction endonuclease. All Uniprot IDs and more information listed in Supplementary Table S1.

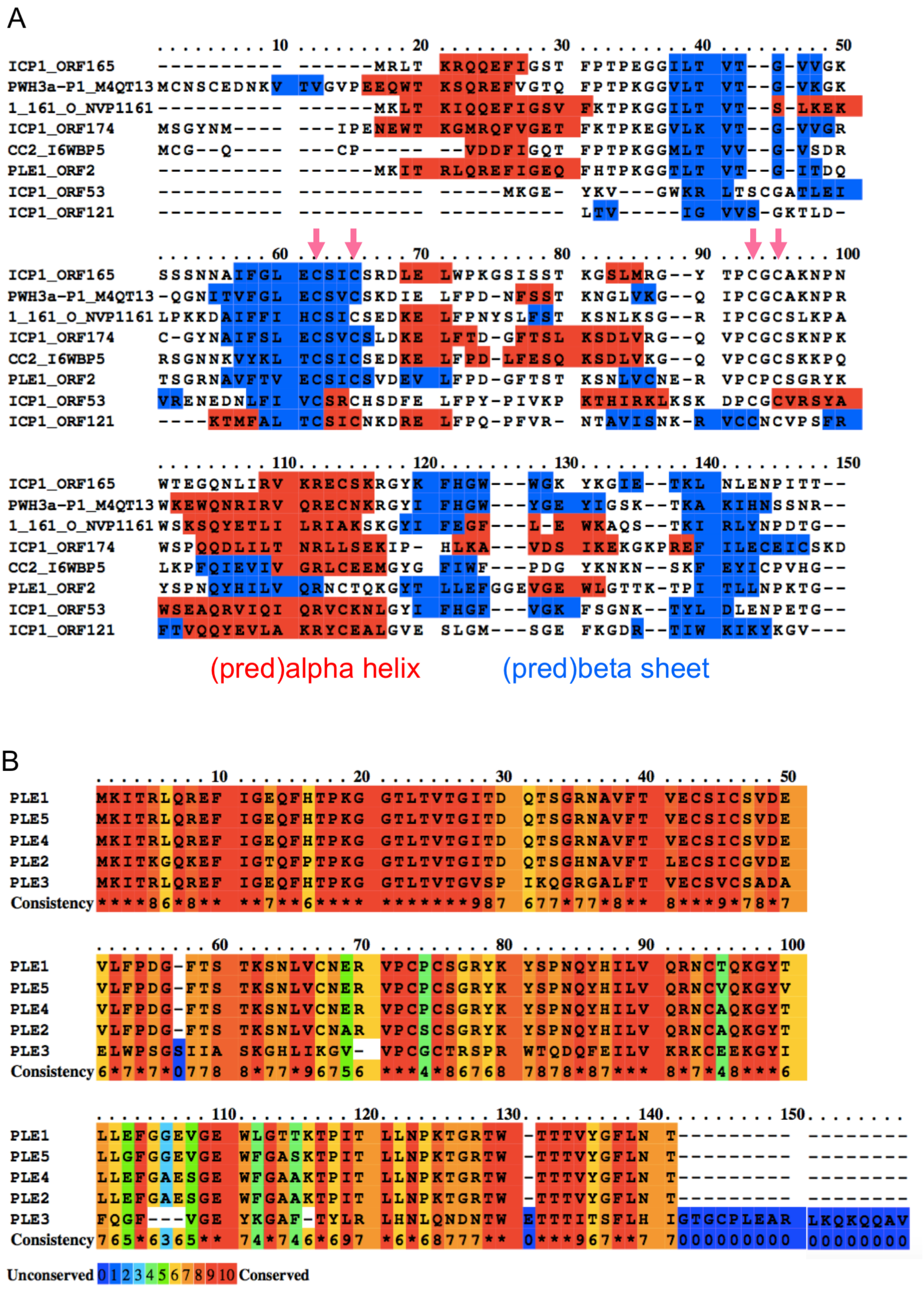
A
(pred)alpha helix
(pred)beta sheet
B
Unconserved
Conserved

**Supplementary Figure 8. Multiple protein sequence alignments**. (**A**) Trimmed multiple protein sequence alignment of N-terminal domains with ~30-50% homology to CapR. Regions highlighted in red are predicted alpha helical structural motifs and regions highlighted in blue are predicted beta sheets. Pink arrows denote conserved cysteine sets. PLE1_ORF2 denotes CapR (**B**) Multiple protein sequence alignment of the CapR from each of the five previously-described PLEs. Conservation indicated by color scale below.

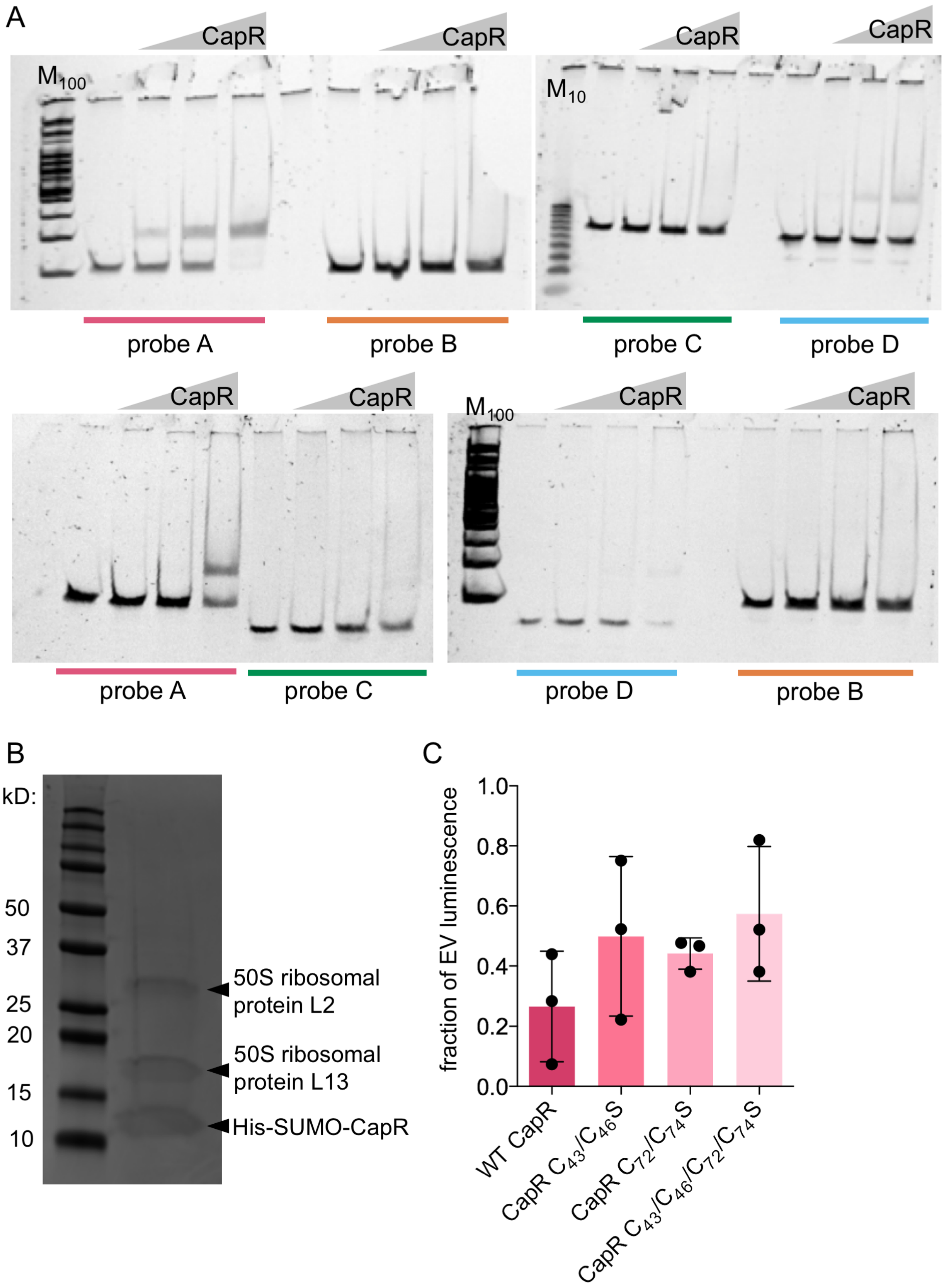
A
CapR
CapR
CapR
CapR
$M_{100}$
$M_{10}$
probe A
probe B
probe C
probe D
CapR
CapR
CapR
CapR
$M_{100}$
probe A
probe C
probe D
probe B
B
kD:
50
37
25
20
15
10
50S ribosomal protein L2
50S ribosomal protein L13
His-SUMO-CapR
C
fraction of EV luminescence
1.0
0.8
0.6
0.4
0.2
0.0
WT CapR
CapR $C_{43}/C_{46}S$
CapR $C_{72}/C_{74}S$
CapR $C_{43}/C_{46}/C_{72}/C_{74}S$

**Supplementary Figure 9. CapR protein activity continued.** (**A**) Biological replicates of EMSA experiment shown in Fig. 6. (**B**) Purified CapR protein. kD indicates molecular weight of bands in the protein standard in the first lane. Protein contaminants identified by mass spectrometry on total elution and matching high-frequency hits to sizes on gel. (**C**) ICP1 capsid NanoLuc reporter expression 16 minutes post-infection of *V. cholerae* hosts expressing wild-type or mutant CapR alleles. Expressed as a fraction of the RLU from infection of an empty vector (EV) control.
